# Effects of forest management on the phenology of early-flowering understory herbs

**DOI:** 10.1101/733907

**Authors:** Franziska M. Willems, J.F. Scheepens, Christian Ammer, Svenja Block, Anna Bucharova, Peter Schall, Melissa Sehrt, Oliver Bossdorf

**Author notes:** **Correspondence:** Franziska Merle Willems.

## Abstract

Many organisms respond to anthropogenic environmental change through shifts in their phenology. In plants, flowering is largely driven by temperature, and therefore affected by climate change. However, on smaller scales climatic conditions are also influenced by other factors, including habitat structure. A group of plants with a particularly distinct phenology are the understorey herbs in temperate forests. In these forests, management alters tree species composition and stand structure and, as a consequence, light conditions and microclimate. Forest management should thus also affect the phenology of understorey herbs. To test this, we recorded the flowering phenology of 20 early-flowering herbs on 100 forest plots varying in management intensity, from near-natural to intensely managed forests, in Central and Southern Germany. We found that in forest stands with a high management intensity the plants flowered on average about two weeks later than in unmanaged forests. This was largely because management also affected microclimate (e.g. spring temperatures of 5.9 °C in managed coniferous, 6.7 in managed deciduous and 7.0 °C in unmanaged deciduous plots), which in turn affected phenology, with plants flowering later on colder and moister forest stands (+4.5 days per −1°C and 2.7 days per 10 % humidity increase). Among forest characteristics, the main tree species as well as the age, overall crown projection area, structural complexity and spatial distribution of trees had the greatest influence on microclimate. Our study demonstrates that forest management alters plant phenology, with potential far-reaching consequences for the ecology and evolution of understorey communities. More generally, our study suggests that besides climate change other drivers of environmental change, too, can influence the phenology of organisms.

## Introduction

Phenology is the study of the timing of recurrent biological events, the biotic and abiotic drivers of this timing, and its variation within and among species (Lieth, 1974). It includes the seasonal timing of key life events, such as animal migration or reproduction, or the leaf-out, flowering and fruiting of plants, which are important for individual fitness. In plants, many phenological events are triggered by biotic and abiotic environmental factors, especially temperature, and are therefore sensitive to climate change (Schwartz, Ahas, & Aasa, 2006; Tang et al., 2016). Long-term observational studies have found earlier leaf-out and changes in the start of flowering associated with climate change across the world (Fitter and Fitter, 2002; Schwartz et al., 2006). Spring-flowering plants seem to be particularly responsive to climate change and often show the largest phenological shifts (Chmielewski, Müller, & Bruns, 2004; Fitter and Fitter, 2002; Renner & Zohner, 2018).

Plants play a key role in many ecosystems, and they interact with many other species. Therefore shifts in plant phenology can have significant consequences for pollinators, food webs, agricultural yields, as well as many ecosystem functions and services such as productivity and carbon cycling (Chmielewski et al.,2004; Cleland et al., 2007; Reilly et al., 1996; Tang et al., 2016;). Understanding the drivers of phenology variation is thus important to predict future states of species abundance and distribution, biogeochemistry and ecosystem productivity, as well as ecosystem services such as pollination (Chuine, 2010; Durant et al., 2005; Høye, Post, Eric, Schmidt, Trøjelsgaard, & Forchhammer, 2013; Kharouba et al., 2018; McKinney et al., 2012; Memmott, Craze, Waser, & Price, 2007; Richardson et al., 2010), and it should also help to inform environmental conservation (Cerdeira Morellato et al., 2016) and to develop adaptive management strategies in a changing world Bellard, Bertelsmeier, Leadley, Thuiller, & Courchamp, 2012; Enquist, Kellermann, Gerst, & Miller-Rushing, 2014; Pacifici et al., 2015; Walther, 2010).

However, our mechanistic understanding of the impact of environmental change on plant phenology is still limited (Richardson et al., 2012). In particular, besides climate warming the influences of other global change drivers – such as land use change – on plant phenology have received little attention. On small scales, climatic conditions such as temperature and humidity are also influenced by a variety of habitat factors such as topography or forest cover (Geiger, Aron and Todhunter, 2003). As a consequence, microclimates can differ from regional climate patterns and affect the timing of phenological events on small spatial scales (Hwang et al., 2011; Ward, Schulze, and Roy, 2018). In forests, stand structure affects the microclimate and light availability (Baker et al., 2014; Chen et al., 1999) and is thus likely to impact flowering phenology of understory herbs. Forest stand structure can be defined as the distribution of trees in space and their variability in size, arrangement, consistency and time (Schall et al., 2018). Stand structure can, for example,be characterized by the main tree species, the ages of trees, their mean diameters at breast height, the basal area covered, or their crown projection area (Schall et al., 2018). Furthermore, stand structural complexity indices (SCI, see for example Zenner and Hibbs (2000)) can combinate several structural attributes (Gossner et al., 2014; del Río et al., 2016) or take the spatial distribution of trees into account (Ehbrecht, Schall, Ammer and Seidel 2017; Penttinen, Stoyan, and Henttonen, 1992).

Changes in forest management alter stand structure in temperate forests and, as a consequence, microclimate conditions. While thinnings and selection cuttings lead to only small increases of radiation at the forest floor (Aussenac 2000; Hale, 2003), clear-cuttings result in drastic and long-persisting changes of the microclimate. In deciduous forests, there is a time window during spring when the leaf-out of trees is not yet completed that allows early spring-flowering species to take full advantage of the available sunlight, moisture and nutrients of the forest floor (Lapointe, 2001). Planting of evergreen coniferous trees – such as Norway spruce (*Picea abies* (L.) H. Karst), one of the most economically important tree species in Europe (Spiecker, 2003) – reduces the light availability during early spring and changes microclimatic conditions. More generelly, all management changes that alter tree species composition and stand structure are likely to also affect the phenology of forest understory herbs, through changes in radiation, microclimate, or other factors. Because of their narrow and distinct flowering period, spring-flowering forest herbs should thus be particularly susceptible to management changes, and therefore they are a particularly relevant study system for exploring forest management effects on plant phenology.

Here, we studied the phenology of 20 early-flowering forest herbs on 100 forest plots of different management type and intensity. We hypothesized that forest management would change forest structure and, as a consequence, microclimatic conditions that impact flowering phenology. We obtained detailed phenology, forest structure and microclimate data for each of the studied plots, and we used structural equation modelling to understand the underlying causal relations, and to disentangle the direct and indirect effects that different forest characteristics and microclimatic variables have on plant phenology. Specifically, we asked the following questions: (i) Does forest management intensity affect plant phenology? (ii) Which forest characteristics are the strongest drivers of phenological variation? And (iii) to what extent do forest characteristics affect phenology directly versus indirectly through changing microclimatic conditions?

## Methods

### Study system

Most forests in Central Europe have a rather low tree diversity and are dominated by only few decicuous tree species (Schulze et al., 2016). Therefore, variation in stand structures is to a substantial degree related to the effects of forest management (Schall et al., 2018). Here we studied the forest plots of the Biodiversity Exploratories project (www.biodiversity-exploratories.de) in Germany, a large-scale platform for ecological research that includes a broad range of forests plots of different management types and intensities (Fischer et al. 2010). We focused on 100 forest plots (100 × 100 m) located in equal parts in two of the three regions of the Biodiversity Exploratories, the Schwäbische Alb in Southwest Germany (long: 9.39°, lat: 48.44°) and the Hainich-Dün in Central Germany (long: 10.47°, lat: 51.16°). The elevation a.s.l. range from 285–550 m in the Hainich-Dün area to 460–860 m on the Schwäbische Alb. Further details on the characteristics of the regions are provided in Fischer et al. (2010). The forests in the study areas are dominated by native deciduous trees, mainly European beech (*Fagus sylvatica* L.). However, some forests were converted to plantations of Norway spruce (*Picea abies*), a coniferous species originally restricted to montane regions, but cultivated for timber in the lowlands since 250 years (see Figure 1 & Schall et al., 2018). The studied plots included (more intensively managed) even-aged deciduous forests at a range of developmental stages, but also uneven-aged and unmanaged deciduous plots, as well as managed even-aged stands of coniferous spruce forests of different age-classes (see Table S6 & Schall et al., 2018). On all but three plots the main tree species was either beech or spruce, and we therefore grouped all beech plots together with the three other hardwood-dominated plots as deciduous forest plots (*N* = 83 plots) for the subsequent analyses, whereas plots dominated by Norway spruce were labelled as coniferous forest plots (*N* = 17 plots).

**Figure 1:**
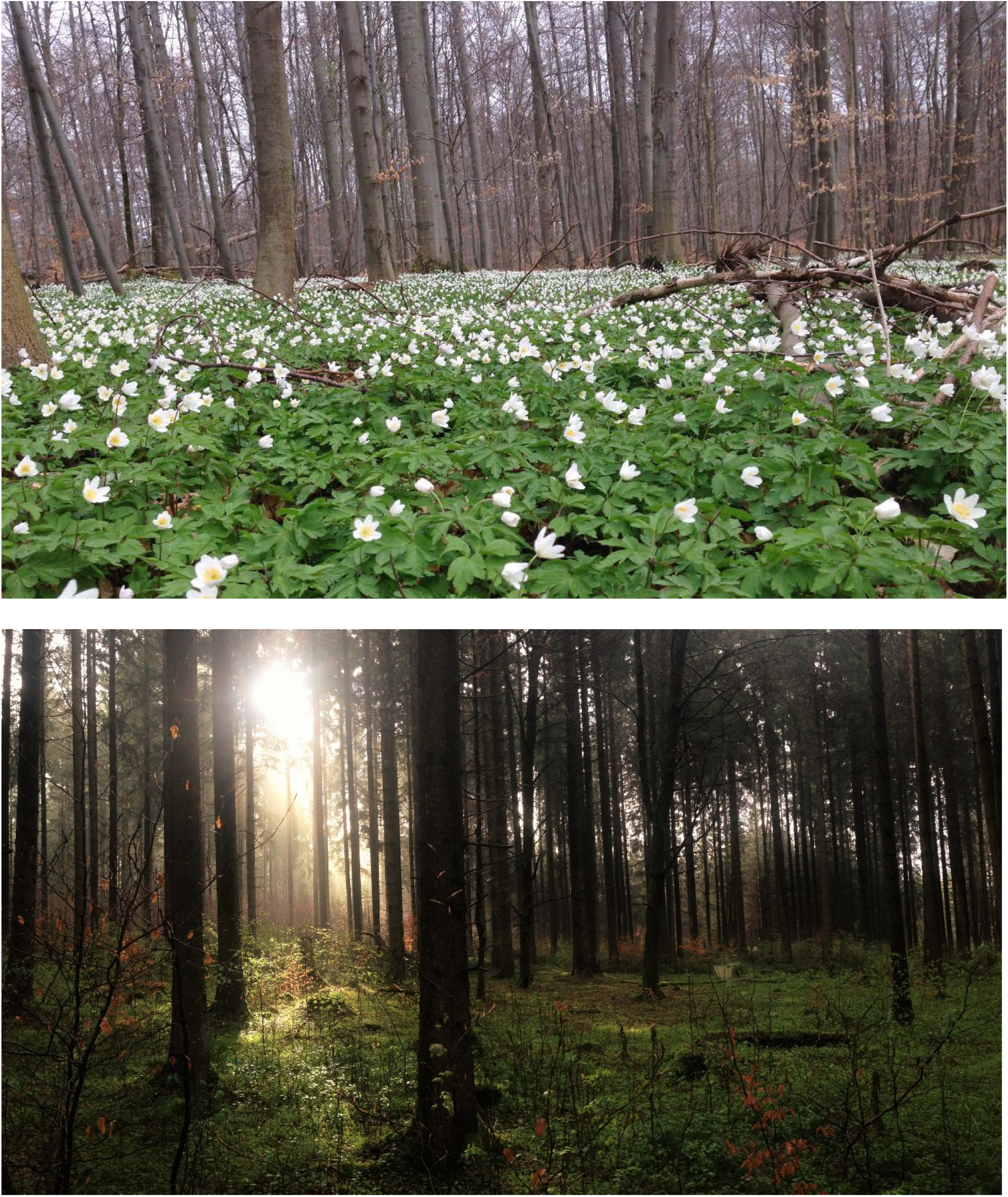
Impressions from the study plots. Top: beech plot at the Hainich-Dün. Bottom: spruce plot at the Schwäbische Alb. Both photos were made in April 2017.

### Phenological monitoring

From March to June 2017, we monitored the phenology of 20 early-flowering herbs in the understories of our study plots (see Figure 2 and Table S1). The monitored species included all common spring-flowering herbs in the plots. We visited all 100 forest plots once per week and monitored the phenology of all plants within a 3 m wide strip outside the 20 × 20 m core area of each plot, corresponding to an area of 224 m^2^ within each 1 ha plot. For each species in each plot, we recorded flowering start as the day of the year with the first fully open flower, and flowering end as the time when no fully open flowers could be found anymore. To be able to determine flowering peaks, we counted the number of open inflorescences or, if plants were abundant on a plot, we estimated the percentages of flowering individuals. We then defined the day of the year with the highest number or percentage of open inflorescences as the day of flowering peak. If there were two days with equal maximum flowering, we used their median as the time of peak flowering. If we visited a plot, and it was apparent that a start, peak or end of flowering had been between the present and past visit, we dated this record back to the previous Monday or Thursday, resulting in an effective half-weekly resolution of our data.

**Figure 2:**
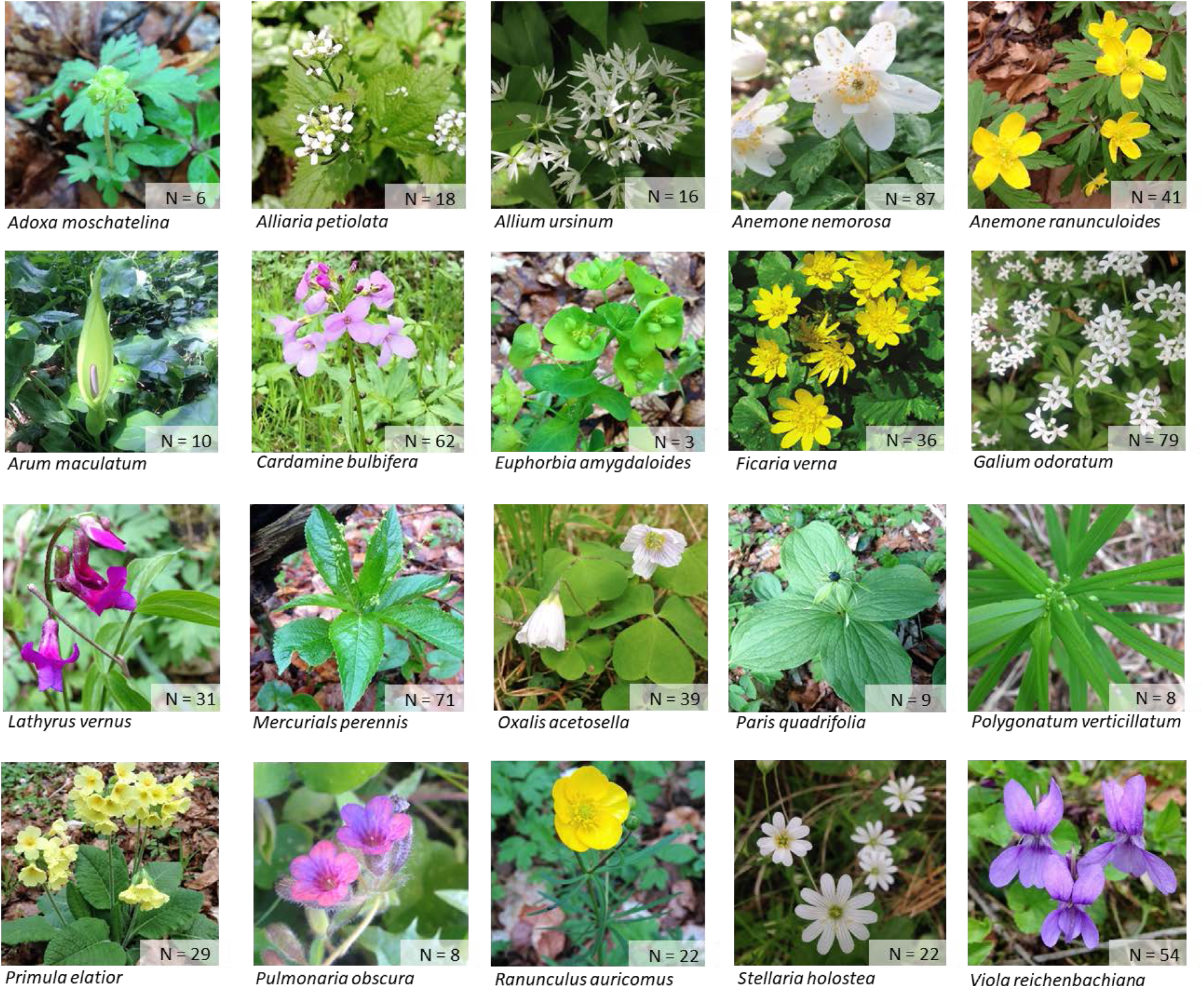
The 20 early-flowering forest understorey species included in our phenology monitoring. For each species, the number of plots with flowering individuals is indicated in the bottom right corner.

### Forest characteristics

The structure of the studied forests is strongly influenced by management, and it can be characterized by differing forest attributes. The required data have been collected in two forest inventories that were conducted on the forest plots of the Biodiversity Exploratories at single-tree level for all living trees with a diameter at breast height ≥ 7 cm. We generally used the data from the most recent inventory (2014-2016) except for 12 plots where we these data were incomplete and we therefore used information from the previous inventory in 2008-2011. Specifically, we used the following variables: main tree species (deciduous vs. coniferous), the mean age of the main tree species, the richness and diversity (inverse Simpson’s index) of tree species, crown projection area, the share of conifers based on crown projection, stand density, the mean diameter at breast height and its standard deviation, and the basal area covered with trees. Furthermore, we used Morisita’s index of dispersion as well as Clapham’s variance mean ratio as measures of horizontal heterogeneity (for both <1: regular, >1: clumping, 1: random; 20 m × 20 m raster cells), and Zenner’s Structural Complexity Index based on tree height as a proxy for vertical structural complexity (Zenner, 1998). We selected these variables because they characterize stand structure, and we expected them to have an influence on microclimatic conditions as well as on light availability and other abiotic and biotic factors. In addition to these individual forest variables, we also tested an index for silvicultural management intensity developed by Schall and Ammer (2013) which combines tree species, stand age and aboveground living and dead wood biomass as three main characteristics of a forest stand into an overall measure of forest management intensity. For an overview of all explanatory variables, see Figure S8.

### Microclimate and other environmental data

Besides the data on forest structure, there is detailed information on local microclimate available for all plots in the Biodiversity Exploratories (Fischer et al., 2010). To be able to test for relationships between microclimate, forest management and phenology, we compiled data for two different potentially relevant time periods, the spring months during which our phenology monitoring took place, and the preceding winter months. For the spring months (February–May 2017), we calculated the average mean air temperature (measured at 10 cm and 2 m height), the growing days (=days with mean temperatures between 10°C to 30°C), the growth sum (= sum of mean day temperatures > 5°C (minus 5)), the warm sum (= sum of mean day temperatures with > 10 °C (minus 10)), mean relative air humidity (measured at 2 m), as well as mean soil moisture and soil temperature (both measured at 10 cm depth). For the winter months (October 2016-January 2017), we also calculated the mean air temperature (measured at 2 m height and 10 cm height), the number of cold days (= days with a temperature minimum < 0°C), the cold sum (= sum of mean day temperatures < 0 °C), the number of cool days (= days with a temperature maximum < 10°C), the number of ice days (= days with a temperature maximum < 0°C), mean relative air humidity (measured at 2 m), as well as mean soil moisture and soil temperature (both measured at 10 cm depth).

In addition to the microclimate data, we also included several geographical variables that we expected to influence abiotic conditions at the stand level, such as exploratory (region), slope (in degrees; average over the plot area) and aspect (= circular average over the plot area, 360° = 0° denote a north facing orientation, 180° indicates a south facing orientation etc.). We combined the data on aspect and slope combined into an aspect×slope variable by multiplying inclination by 1 for south-, −1 for north-, and 0.5 for east- and west-facing slopes, to be able to distinguish slopes in the four cardinal directions which are known to differ in their microclimatic conditions (Dahlgren, Zeipel, and Ehrlén, 2007). Elevation above sea level is confounded with region and therefore not included as an explanatory variable. For an overview of all explanatory variables, see Figure S8.

### Data analysis

Our data analyses following a two-step logic. First, we used univariate linear regression to test the effects of forest management intensity, as well as individual forest characteristics and microclimatic variables on flowering time for all species separately. Second, we selected a subset of these variables for structural equation modelling, to understand the relationships between forest characteristics and microclimate, and disentangle direct and indirect effects on plant phenology. Prior to the data analyses, we checked all variables for outliers, and if outliers clearly resulted from measurement errors, we removed them from our data set. Moreover, for the statistical analyses we excluded two species that were flowering on less than eight plots (Table S1), *Adoxa moschatellina* L. and *Euphorbia amygdaloides* L.

Using linear regression analyses, we calculated *R*^*2*^-values, standardized regression coefficients and *P*-values (corrected for multiply testing using FDR) for each the relationships between each forest trait and microclimatic variable and the phenology of each studied species. We used these results to make an informed preselection of variables for the subsequent structural equation model (see next section), since especially the microclimatic variables included several temperature proxies with high levels of collinearity. All data analyses were conducted using R (R Core Team, 2018). Standardized regression coefficients were derived using the “QuantPsyc” package (Fletcher, 2012).

Next, we conducted confirmatory path analysis over all species based on piecewise fitting of component hierarchical linear mixed-effects models (Lefcheck, 2016; Shipley, 2009). Path analysis or structural equation modelling is a powerful, multivariate technique used increasingly in ecology to evaluate complex multivariate causal relationships, particularly with observational data that often includes substantial collinearity. Structural equation models (SEMs) differ from many other modelling approaches as they test the direct and indirect effects in pre-assumed causal relationships (Fan et al. 2016). In our analysis we used the “piecewiseSEM” package (Lefcheck, 2016). In piecewise SEM, each set of relationships is estimated independently (or ‘locally’). For each response variable, the process decomposes the network into the corresponding simple or multiple linear regressions, which are evaluated separately, and then re-combined afterwards to draw conclusions about the full model (Lefcheck, 2016). The relationships between variables can then be visualized through path diagrams where arrows denote which variables are influencing (and are influenced by) other variables.

Prior to our path analyses we checked for additivity and linearity of individual variables. We used correlation matrices (Figures S2) and variance inflation factors (with a cut-off value of 4) to check for collinearity among the explanatory variables, to avoid inclusion of highly correlated variables. We used simple regression plots to confirm linearity. Furthermore, to check the statistical assumptions of linear models – normality and homogeneity of residuals – we visually inspected histograms of the standardized residuals, Q-Q-Plots and residual scatter plots, as well as calculations of skewness and kurtosis. The skewness and kurtosis values were all within the guidelines set by Kline (2015) and also below the more conservative threshold set by Ryu (2011).

The subset of forest characteristics that we included in the SEM, after checking for collinearity, were: crown projection area, variance mean ratio, structural complexity index, diameter at breast height, its standard deviation, the percentage of coniferous trees. Diameter at breast height was selected as an explanatory variable over age and density because it was the best proxy for the developmental stage of a forest. After the exclusion of highly correlated variables and based on the simple linear regressions results (considering average *r*^*2*^-values and standardized regression coefficients), mean spring air temperature and spring relative humidity were the only microclimatic variables we included in the SEM. Because other geographical or environmental factors might also influence plant phenology, we additionally included aspect×slope as well as exploratory (region) as explanatory variables in the SEM. In the sub-model with flowering peak as a response variable and forest characteristics and microclimatic variables as explanatory variables, we included species identity as a random variable. To test whether the forest characteristics influence the local microclimate, we set both spring air temperature and spring relative humidity also as response variables, while using the forest characteristics as well as other geographical factors as explanatory variables. The complete dataset included 724 data points, but since 122 rows had missing values for at least one of the variables, we analysed the full SEM with 602 data points.

We evaluated the overall path model using Shipley’s test of directed separation (Shipley 2009), which yields a Fisher’s C statistic comparable to a χ^2^. A *P*-value > 0.05 indicates that a model can adequately reproduce the hypothesized causal network. Fisher’s C is then used to calculate the Akaike Information Criterion (AIC), or a corrected AIC for small sample sizes (AICc), to compare model fits. We calculated both marginal and conditional *R*^2^-values, where the former describes the proportion of variance explained by only fixed factors, whereas the latter describes the variance explained by fixed and random factors. Starting with a full model based on *a priori* knowledge of interactions that included all the above-mentioned variables, we used a backwards stepwise elimination process based on AICc to remove non-significant pathways. Additionally, we used d-separation tests to evaluate whether any non-hypothesized independent paths were significant, and whether the models could be improved by including any of the missing paths.

## Results

The onset of flowering in our study species ranged from mid-March (*Mercurialis perennis* L., *Primula elatior* (L.) Hill, *Anemone nemorosa* L.) to the beginning of May (*Galium odoratum* (L.) Scop.), *Arum maculatum* L., *Polygonatum verticillatum* (L.) All.). Similarly, the peak flowering time of the different species ranged from the end of March until the end of May. For some species, the flowering period ended already in mid-May while others continued to flower until mid-June. Besides these species differences in mean onset, peak and end of flowering, we also found large differences among species in their levels of among-plot variation (see Figure 4). Some species had very narrow ranges, e.g. the flowering peak of *Galium odoratum* varied only by 10 days across the 79 studied plots, whereas for *Anemone nemorosa* (*N* = 87) and *Mercurialis perennis* (*N* = 71) the flowering peaks differed by up to 42 and 46 days, respectively. For an overview of mean flowering start, peak and end, as well as the respective *N*, of all species see Supplement Table S1.

**Figure 3:**
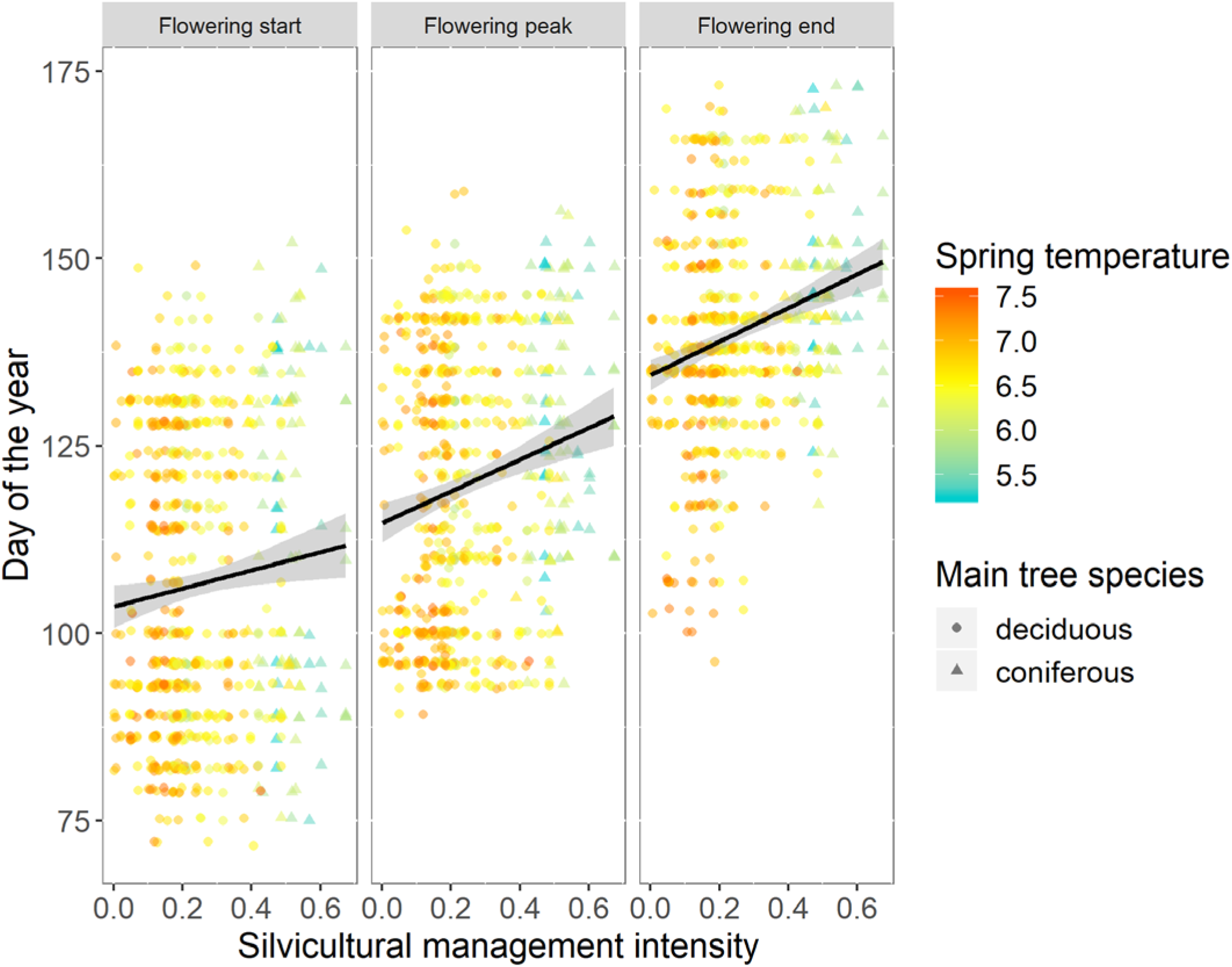
The relationships between silvicultural management intensity and flowering start, peak and end, respectively, across all studied species. Each point represents a plot by species combination. Silvicultural management intensity values can range from 0 (lowest management intensity) to 1 (highest management intensity). The shape and colour of the symbols code for main tree species and mean spring temperature (see legend). The lines are linear regressions with 95% confidence intervals.

**Figure 4:**
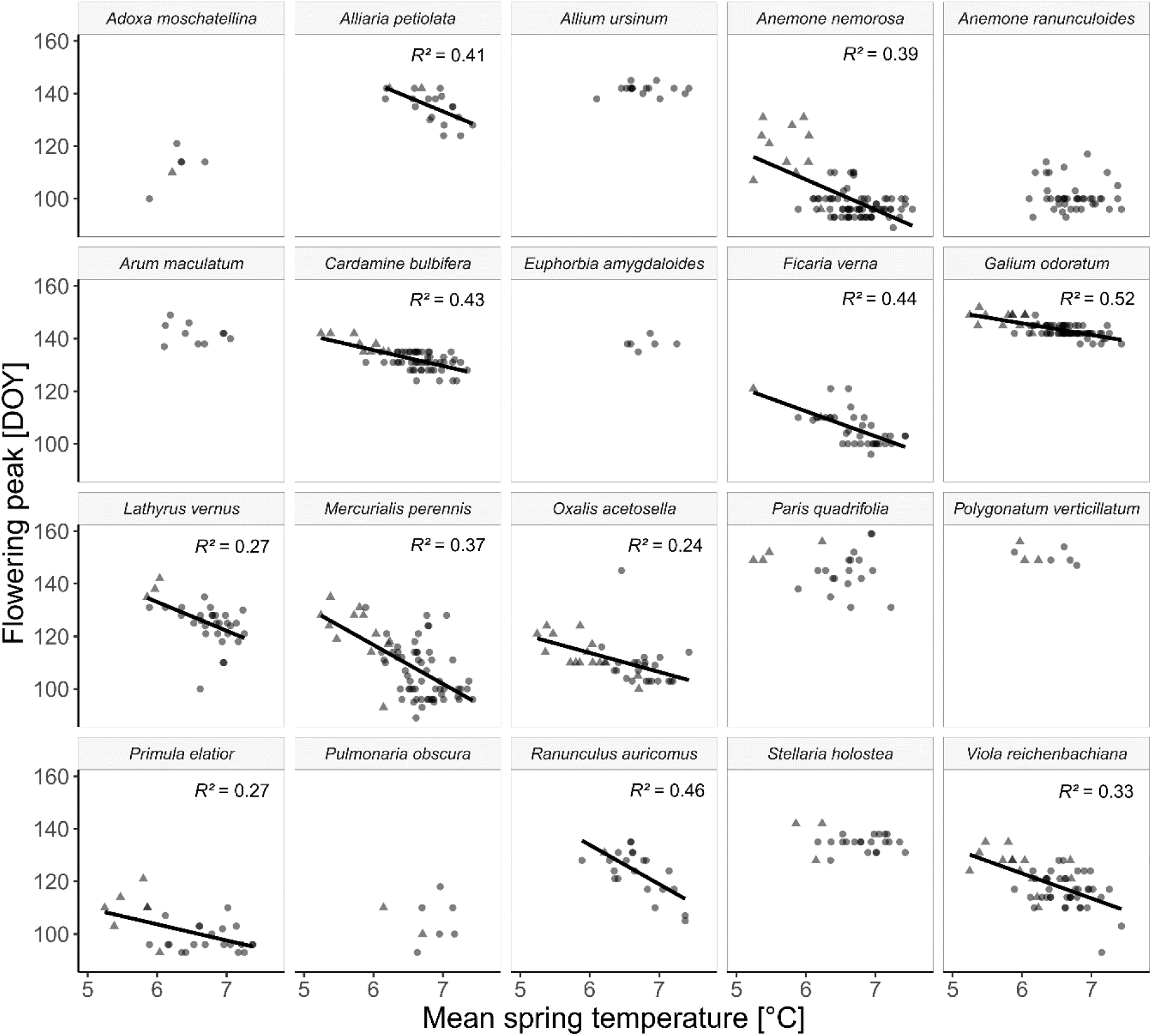
Regression of flowering peak (DOY, day of year) against mean spring temperature. Each point represents a forest plot, and the shape of each point indicates whether the main tree species is deciduous (circle) or coniferous (triangle). For significant regressions, the fitted regression lines are plotted. All regression coefficients are listed in Table S2.

### Impact of forest management on phenology

Across all studied species, forest understory herbs growing on plots with a high silvicultural management intensity had a significantly delayed start, peak and end of their flowering periods (Figure 3, Flowering start: correlation coefficient β = 12.11, adjusted *R*^*2*^ = 0.01, *P*-value = 0.013. Flowering peak: correlation coefficient β = 21.19, adjusted *R*^*2*^ = 0.03, *P*-value < 0.001. Flowering end: correlation coefficient β = 22.43, adjusted *R*^*2*^ = 0.06, *P*-value < 0.001.) On plots with the highest forest management intensity, the average peak of flowering was over two week later than on plots with the lowest management intensity (day of the year 127-129 in managed spruce forest and day of the year 118 in managed beech forests versus day of the year 113 in unmanaged beech forest). Generally, plants flowered later on plots dominated by coniferous trees than on deciduous forest plots (Figure 3 and 4). These general patterns were also reflected at the level of individual species: in all but one of the studied species, there was a positive (albeit not always significant) relationship between silvicultural management intensity and peak flowering (Table 1), with some of the strongest effects observed in *Primula elatior, Anemone nemorosa* and *Galium odoratum*, all of them emblematic spring flowers in temperate forests. For detailed regression results, see Table 1 and Tables S5 and S6.

**Table 1:** Relationships between forest characteristics and microclimate (last two columns), and the peak flowering of different plant species. The values are standardized regression coefficients derived from linear regressions of flowering peak against the different forest trait and microclimate variables, with significant values in bold (corrected using FDR). Blue colours indicate positive relationships, orange colours indicate negative relationships, and colour intensity is proportional to effect size. Age = mean age of main tree species, AspSlope = a combination of inclination and exposition, BA = basal area covered by trees, CPA = crown projection area, Con = Coniferous, dbh = diameter at breast height, dbh SD = standard deviation of dbh, density = stand density, Div = species richness of trees, Div 2D = inverse Simpson’s index for trees, Moristia = Morisita’s index of dispersion, SCI = Zenner’s structural complexity index, SMI = silvicultural management intensity, VMR= Clapham’s variance mean ratio. Ta = air temperature and rH = relative humidity, with both climate variables measured at 200 cm height during February – May 2017. See Table S8 for a more detailed explanation of the different explanatory variables, and Tables S3-S6 for the corresponding *R*^2^ values and unstandardized regression coefficients for all regressions.

|  | Age | AspSlope | BA | Con BA | Con CPA | CPA | Dbh | Dbh SD | Density | Div | Div 2D | Morisita | SCI | SMI | VMR | Ta | rH |
| --- | --- | --- | --- | --- | --- | --- | --- | --- | --- | --- | --- | --- | --- | --- | --- | --- | --- |
| *Alliaria petiolata* | -0.41 | -0.08 | 0.10 | 0.42 | 0.41 | **0.06** | -0.04 | -0.27 | 0.23 | 0.32 | 0.26 | -0.19 | -0.27 | 0.41 | -0.16 | **-0.64** | 0.26 |
| *Allium ursinum* | -0.52 | 0.13 | -0.43 |  |  | **-0.39** | -0.10 | -0.29 | 0.19 | 0.24 | 0.55 | -0.14 | -0.33 | 0.26 | 0.15 | 0.06 | -0.17 |
| *Anemone nemorosa* | **-0.37** | 0.01 | **0.46** | **0.82** | **0.82** | 0.08 | 0.16 | -0.22 | -0.04 | -0.14 | -0.17 | -0.13 | **-0.34** | **0.54** | -0.13 | **-0.63** | **0.37** |
| *Anemone ranunculoides* | -0.11 | **-0.41** | 0.00 | 0.32 | 0.34 | **0.08** | -0.19 | -0.20 | 0.28 | 0.20 | 0.03 | 0.08 | -0.07 | 0.07 | 0.22 | -0.11 | 0.01 |
| *Arum maculatum* | -0.55 | **-0.63** | -0.10 | 0.19 | 0.18 | **0.06** | -0.36 | -0.35 | 0.57 | 0.11 | 0.06 | 0.06 | 0.22 | 0.22 | -0.12 | -0.29 | 0.07 |
| *Cardamine bulbifera* | -0.22 | **-0.33** | **0.48** | **0.58** | **0.59** | 0.11 | 0.31 | -0.01 | -0.22 | -0.31 | **-0.35** | **-0.35** | -0.21 | 0.32 | **-0.37** | **-0.65** | **0.56** |
| *Ficaria verna* | -0.31 | -0.02 | 0.01 | 0.37 | 0.38 | **-0.16** | 0.01 | -0.18 | 0.08 | -0.03 | -0.06 | -0.18 | -0.36 | **0.45** | -0.16 | **-0.67** | 0.34 |
| *Galium odoratum* | **-0.39** | -0.24 | **0.36** | **0.60** | **0.61** | 0.08 | 0.14 | -0.19 | 0.03 | -0.21 | -0.21 | -0.17 | **-0.42** | **0.49** | -0.19 | **-0.72** | **0.36** |
| *Lathyrus vernus* | -0.14 | **-0.28** | 0.13 | **0.50** | **0.51** | -0.28 | 0.09 | -0.17 | -0.22 | -0.28 | -0.40 | -0.10 | -0.37 | 0.42 | 0.12 | **-0.52** | **0.53** |
| *Mercurialis perennis* | **-0.39** | -0.23 | 0.19 | **0.53** | **0.53** | -0.08 | 0.07 | -0.27 | 0.02 | -0.06 | -0.08 | -0.22 | -0.32 | **0.48** | -0.15 | **-0.61** | **0.45** |
| *Oxalis acetosella* | -0.16 | **0.12** | 0.05 | 0.28 | 0.29 | **-0.24** | 0.02 | -0.08 | 0.03 | -0.21 | -0.13 | -0.13 | -0.18 | 0.30 | -0.11 | **-0.49** | 0.38 |
| *Paris quadrifolia* | -0.14 | 0.49 | 0.19 | 0.45 | 0.43 | -0.14 | 0.24 | -0.06 | -0.12 | 0.24 | 0.04 | 0.02 | -0.40 | 0.41 | 0.02 | -0.12 | -0.07 |
| *Polygonatum verticillatum* | -0.21 | **0.31** | 0.10 | 0.15 | 0.16 | **-0.03** | -0.07 | **-0.40** | 0.09 | -0.45 | -0.20 | 0.04 | -0.34 | 0.10 | -0.04 | -0.45 | 0.69 |
| *Primula elatior* | -0.26 | -0.09 | 0.46 | **0.63** | **0.63** | -0.04 | 0.24 | -0.10 | -0.09 | -0.25 | -0.19 | -0.13 | -0.21 | **0.57** | -0.11 | **-0.52** | **0.63** |
| *Pulmonaria obscura* | 0.45 | 0.04 | 0.23 | -0.06 | -0.08 | 0.19 | 0.35 | 0.46 | -0.37 | -0.27 | 0.07 | -0.28 | 0.79 | -0.46 | -0.36 | 0.00 | -0.32 |
| *Ranunculus auricomus* | -0.29 | 0.24 | -0.25 | 0.23 | 0.23 | -0.36 | 0.11 | -0.20 | -0.10 | -0.22 | -0.24 | -0.25 | -0.32 | 0.26 | -0.15 | **-0.68** | 0.06 |
| *Stellaria holostea* | -0.38 | -0.42 | 0.07 | 0.32 | 0.35 | **0.04** | -0.23 | **-0.42** | 0.21 | 0.27 | 0.05 | 0.08 | -0.19 | 0.28 | 0.16 | -0.14 | 0.07 |
| *Viola reichenbachiana* | -0.24 | -0.28 | **0.45** | **0.51** | **0.52** | 0.08 | 0.31 | 0.01 | -0.24 | **-0.36** | **-0.41** | -0.24 | -0.20 | 0.31 | -0.33 | **-0.57** | **0.50** |
| Average across all species | -0.26 | -0.09 | 0.14 | 0.40 | 0.41 | -0.05 | 0.06 | -0.16 | 0.02 | -0.08 | -0.08 | -0.12 | -0.19 | 0.30 | -0.10 | -0.43 | 0.26 |

Since forest management affects many aspects of forest structure simultaneously (see Table S7), we used linear regressions to understand which specific forest characteristics were strongly related to variation in plant phenology. We found the strongest statistical associations with flowering peak for the percentage of the crown projection area and the basal area that is taken up by coniferous trees (with an average standardized correlation coefficient of 0.41 and 0.40, and mean *R*^*2*^ = 0.20 and 0.21, maximum *R*^*2*^ = 0.67, respectively) (Table 1). The higher the percentage of coniferous trees was, the later the understory herbs tended to flower (see also Figure 3). Furthermore, plants flowered later in younger forest stands (average standardized correlation coefficient −0.26, with a mean *R*^*2*^ = 0.12, maximum *R*^*2*^ = 0.30) and those with a low structural complexity (average standardized correlation coefficient = −0.19, with a mean *R*^*2*^ = 0.15 and a maximum *R*^*2*^ = 0.89). Table 1 gives an overview of the standardized regression coefficients of all forest characteristics, and the corresponding *R*^*2*^ values and unstandardized regression coefficients are provided in Table S4 and S5.

### Impact of microclimate on phenology

We found that microclimatic conditions varied substantially between different forest plots, and that this was partly related to forest management (Figure 3). For instance, on managed forest plots the mean spring temperatures were substantially lower than on unmanaged forest plots (5.9 °C on managed coniferous, 6.7 on managed deciduous and 7.0 °C on unmanaged deciduous plots). The temperature differences were significant (*P* < 0.001) and the pattern is the same among forest plots within each region (Table S7). Microclimate, in turn, was significantly correlated with plant phenology. Higher spring and winter temperatures were generally associated with earlier flowering, whereas higher humidity was correlated with later flowering (Tables 1 and S2, Figures 4 and S1). Of all microclimatic variables, mean spring temperature measured at 2 m height explained most of the variability of the peak flowering across all species (mean *R*^2^ = 0.25, maximum *R*^2^ = 0.52, for *R*^2^ values of all linear regression see: Supplement Table S3). On average, plants flowered 4.5 days earlier per 1 °C temperature increase, and all plants except *Allium ursinum* L. and *Pulmonaria obscura* Dumont. flowered earlier on plots with higher mean spring temperatures. However, the magnitudes of the responses varied substantially among species, ranging from a change of over 12 days per 1 °C for *Mercurialis perennis* to only minor changes in flowering time of around 1 day per 1 °C for *Paris quadrifolia* L.

For a comparison of all standardized and unstandardized regression coefficients of all microclimatic variables see Tables S2 and S4. Of all moisture related variables, relative humidity during spring was the best predictor of peak flowering (mean *R*^*2*^ = 0.15, maximum *R*^*2*^ = 0.47, see Tables S2-S4 and Figure S1) and was therefore included in the SEM. On average, plants flowered 2.7 days later per 10 % increase of relative humidity.

### Interactions among forest management, microclimate and phenology

The piecewise SEM confirms that, on average, plants flowered earlier on warmer less humid plots and it is a good fit to the data (Fisher’s C = 8.364, df = 12, *P* = 0.756, see Figure 5 and Table 2). It also shows that most of the forest characteristics – percentage of coniferous trees, crown projection area, variance mean ratio and structural complexity index – had a significant influence on the forest microclimate. In particular, spring temperatures were lower on coniferous forest plots than on deciduous forest plots, and forest plots with a lower crown projection area (reflecting forest age), variance mean ratio (reflecting horizontal heterogeneity) and structural complexity were also colder than older and more heterogeneous and structurally complex forest plots. The relative humidity was higher in forest stands with a higher percentage of coniferous trees, and it was lower on warmer plots. Plots located in the Hainich region were generally warmer and more humid and plants tended to flower earlier there than on the Schwäbische Alb (Figure 5 and Table 2). Furthermore, a high percentage of coniferous trees had an equally strong direct effect on the timing of flowering peak, with plants growing on forest stands dominated by Norway spruce flowering later than those in deciduous forests. All unstandardized and standardized estimates of the path coefficients, their degrees of freedom, standard errors, critical values and *P*-values are listed in Table 2.

**Table 2:** Unstandardized and standardized estimates of the path coefficients derived from the piecewise SEM, with their respective standard errors (SE), degrees of freedom (DF), critical- and *P*-values. Only significant paths are listed. CPA = crown projection area, SCI = Zenner’s Structural Complexity Index, VMR = Clapham’s variance-mean ratio, % conifers = percentage of coniferous trees and Exploratory = region.

| Response | Predictor | Estimate | SE | DF | Critical Value | *P*-value | Std. Estimate | Sig. |
| --- | --- | --- | --- | --- | --- | --- | --- | --- |
| Peak flowering | Temperature | -3.52 | 0.91 | 571.00 | -3.88 | 0.000 | -0.09 | *** |
| Peak flowering | Relative humidity | 0.49 | 0.16 | 571.00 | 3.10 | 0.002 | 0.07 | ** |
| Peak flowering | % conifers | 0.0001 | 0.01 | 571.00 | 3.16 | 0.002 | 0.07 | ** |
| Peak flowering | Exploratory (Hainich) | -3.00 | 0.75 | 571.00 | -3.97 | 0.000 | -0.08 | *** |
| Temperature | CPA | 0.00 | 0.00 | 587.00 | 7.54 | 0.000 | 0.23 | *** |
| Temperature | % conifers | -0.01 | 0.00 | 587.00 | -18.96 | 0.000 | -0.64 | *** |
| Temperature | SCI | 0.06 | 0.02 | 587.00 | 2.87 | 0.004 | 0.09 | ** |
| Temperature | VMR | 0.02 | 0.01 | 587.00 | 3.80 | 0.000 | 0.11 | *** |
| Temperature | Exploratory (Hainich) | 0.09 | 0.03 | 587.00 | 2.79 | 0.005 | 0.09 | ** |
| Relative humidity | % conifers | 0.02 | 0.00 | 589.00 | 8.64 | 0.000 | 0.35 | *** |
| Relative humidity | Temperature | -2.89 | 0.20 | 589.00 | -14.26 | 0.000 | -0.58 | *** |
| Relative humidity | Exploratory (Hainich) | 2.68 | 0.16 | 589.00 | 17.13 | 0.000 | 0.55 | *** |

**Figure 5:**
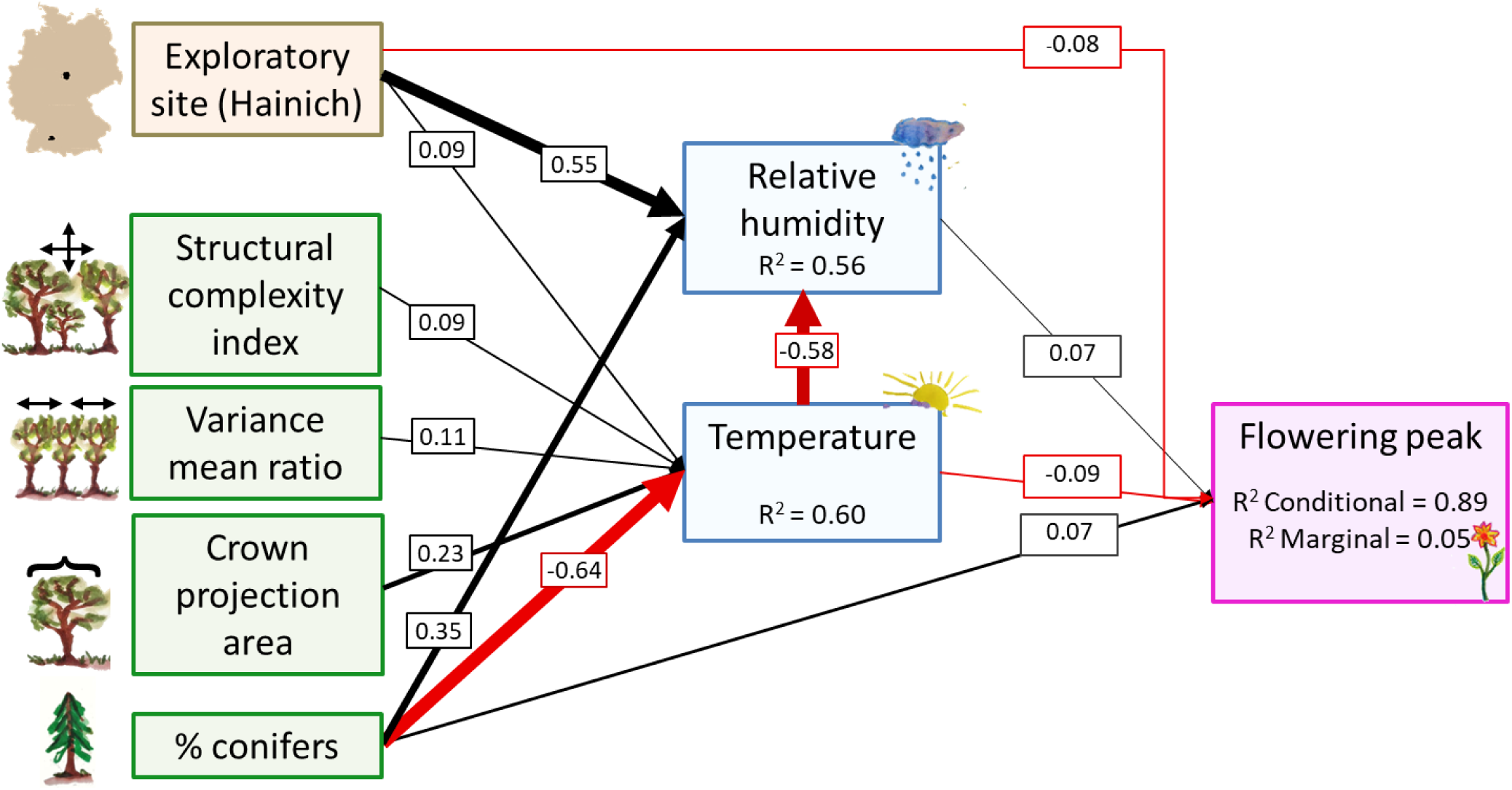
Results of the piecewise structural equation model (SEM) testing for direct and indirect relationships among forest characteristics, geographic parameters, microclimatic variables and the timing of peak flowering of forest understory herbs. Arrows represent unidirectional relationships among variables; only significant paths (*P* < 0.05) are shown. Black arrows are positive relationships, red arrows negatives ones. The thickness of the arrows is proportional to the magnitudes of the standardized regression coefficient, which are also plotted on the arrows. The *R*^2^ values for component models are also given for each response variable. In the model with flowering peak as a response variable, we included the species as random factor. The overall model is a good fit to the data: Fisher’s C = 8.364, *df* = 12, *P-*value = 0.756.

## Discussion

Many organisms respond to anthropogenic environmental change through shifts in their timing of phenological events, and these changes can have far-reaching consequences for the ecology and evolution of ecological communities (Rudolf 2019). It is therefore important to understand the different potential drivers of phenological changes. Here, we disentangled direct and indirect effects that microclimate and forest management have on the phenology of understory herbs. We found that plants flowered later in intensely managed forests than in unmanaged forests. Much of this was because forest management affected microclimate, which in turn affected phenology, with plants flowering later on colder and moister forest stands. Our study thus demonstrates that besides climate change other drivers of environmental change, such as forest management, can influence the phenology of organisms.

### Impact of forest management and forest characteristics on phenology

While climate-related shifts of phenology are widely studied and accepted (e.g Fitter and Fitter, 2002; Parmesan and Yohe, 2003; Wolkovich et al., 2012; Cook, Wolkovich, and Parmesan, 2012), the impacts of other global change drivers, such as land use, have received much less attention. However, land use can also influence life-history traits, such as phenology, and can even cause genetic differentiation in phenological traits (Völler et al., 2013; Völler et al., 2017). Our study demonstrated that understory herbs occurring on forest plots with a high silvicultural management intensity had a significantly delayed start, peak and end of their flowering periods. On forest stands with the highest forest management intensity, the plants flowered on average about two weeks later than those growing in unmanaged forests. Among the different forest characteristics, the percentage of coniferous trees, the age of the trees and the structural complexity of a forest stand were the strongest drivers of phenological variation. Plants generally flowered later on plots dominated by coniferous trees that were relatively young and structurally less complex.

Within the last years, the evidence that land-use change affects the phenology of plants and animals has grown. Zhang, Liu and Herebry (2019) showed that land cover and land use change drive crop phenology, especially in intensively managed agricultural landscapes. Similarly, Altermatt (2012) showed that temperature-related phenological shifts of butterflies depend on their habitat. Leong, Ponisio, Kremen, Thorp and Roderick (2016) found that bee phenology differed between urban and agricultural habitats. For plants it has been suggested that the combined effects of climate change and land-use change accelerated vegetation green-up, with human-managed ecosystems greening up faster than their natural counterparts (Wang et al., 2018).

One might argue that the prolongation of the flowering period through diverse forest management at the landscape scale may improve resource availability and heterogeneity for consumers (such as bees). However, this probability for this is questionable, since abundance of many species (and thus their resource availability) was lower on the intensely managed plots we monitored. Furthermore, within our study regions, planting Norway Spruce is a forest management action, and there are no unmanaged coniferous forests. To tease apart management types from tree species identity, it would be scientifically ideal to compare managed with unmanaged spruce plots, if the latter would exist Therefore, comparing plant phenology also between unmanaged and managed coniferous forest (in other regions) would be a worthwhile focus for future research.

### Impact of microclimate on phenology

Our studied forest plots not only different in their management, but also in their microclimate. Both simple linear regressions and the SEMs confirmed that the flowering phenology of spring-flowering understory herbs was affected by these microclimatic conditions, with higher spring (and winter) temperatures resulting in earlier flowering, and higher relative humidity associated with later flowering. The plants flowered on average 4.5 days earlier per +1 °C temperature difference. This magnitude of change corresponds very well with the response of plants to interannual temperature variation observed in previous studies. For example, Heikinheimo and Lappalainen (1997) suggested that a springtime temperature increase of 1°C can result in flower buds bursting approximately 4 days earlier, based on phenological long-term data for eleven plant taxa (trees, shrubs and forest understory herbs) in Finland. In Britain, the average first flowering of 385 plant species (trees, shrubs and herbs) was advanced by 4.5 days in the 1990s compared to the previous four decades, and in relation to climate the effect size was also 4.3 to 6 days per 1°C increase in mean monthly temperature for spring flowering species (Fitter and Fitter, 2002). Moreover, an analysis of a large phenological network data set showed that across Europe phenological shifts match the warming pattern in Europe (Menzel et al., 2006). Our data show that such climatic differences, and the associated very similar changes in phenology, can also occur on much smaller scales, and that microclimatic patterns can differ substantially from regional climate patterns (Hwang et al., 2011; Ward et al., 2018), and we therefore may need to take them into account when projecting effects of climate change on phenology (De Frenne et al., 2013; Franklin et al. 2013).

We also found that the magnitudes of the temperature-associated phenology changes varied substantially among species. This is consistent with several previous studies. Fitter and Fitter (2002), for example, found that annual plants are more likely to flower earlier than congeneric perennials, and insect-pollinated species more likely than wind-pollinated ones. Such differences in the phenological response might ultimately alter the structures of plant communities. Roberts et al. (2015) predicted that interspecific differences would change the order of spring phenology in temperate forests, which in turn would change hierarchies of light competition and thus potentially the composition of temperate forests. Furthermore, even if the majority of species flower earlier, some may still show insignificant trends or even delayed flowering. In a long-term study of 490 species, Cook et al. (2012) demonstrated that the interaction of fall/winter chilling (i.e. vernalization) and spring warming sensitivities explains much of the apparently paradoxical behaviour of non-responding species, or of species that show delayed spring events despite local warming. That, generally, both warmer spring and winter temperatures are correlated with earlier flowering in our study, indicates that potential vernalization requirements are probably met for (most of) our plants.

High humidity, on average, delayed flowering by around 2.7 days per 10% increase of relative humidity, and the phenological responses of plants to humidity changes were fairly consistent. The findings of previous studies were ambiguous. While some suggested that humidity is crucial for plant phenology (Laube et al., 2014; Matthews and Mazer, 2016), others found no evidence for a significant role of air humidity for plant phenology (Abu-Asab et al., 2001; Zipf and Primack, 2017). Thus, phenological responses to humidity generally seem to be more complex and species-dependent, and they may depend on interactions with other factors.

### Interactions among forest management, microclimate and phenology

The SEM confirmed that microclimatic conditions – spring temperature and relative humidity – are influenced by forest structure which is strongly influenced by forest management. Forest structure generally seems to have a stronger effect on temperature than on relative humidity. Our results confirm those of (Augusto, Dupouey, and Ranger 2003; Nihlgard, 1969), showing that forest dominated by Norway spruce tend to be colder and moister than those dominated by European beech. A particularly interesting result is that less spatially heterogeneous and structurally complex forest plots with a low crown projection area are colder. This may seem couterintuitive at first, because during the day plots with a low crown projection area allow more light to penetrate the canopy and are therefore warmer. However, this trend reverses during the night where plots with a low crown projection area are colder (see Figure S3), presumably due to a sheltering effect of large tree crowns, which reduce convection, mixing of air and infrared reflection (Geiger et al. 2003; von Arx et al., 2013). Since the trend during the night is stronger than during the day the net effect is a cooling under lower crown projection areas. The planting of Norway spruce instead of beech is one of the most critical management decisions. Besides their narrower crown width to diameter ratio to beech, spruce plantations differ from beech forests in many other characteristics such as stand density, size distribution, age, horizontal/spatial- and vertical patterns (Schall et al., 2018). A reason why forest stands dominated by conifers are colder is that particularly in early spring, when deciduous trees have not completed their leaf-out yet, they allow much less light to reach the forest floor and thus do not warm up as much during the day.

In our study, the dominant tree species affected plant phenology not only indirectly, through altering microclimate, but also directly. This direct effect is almost as strong as the effect of temperature, and it must result from other abiotic or biotic factors, besides temperature and humidity, that the dominating tree species in a forest affects. The two most likely candidate factors are light and soil conditions. Evergreen, coniferous trees create much darker conditions on the forest floor during spring, which may be crucial for the development of the understory vegetation (Tinya et al., 2009). Moreover, coniferous forests are also known to differ in various biotic and abiotic traits – many soil properties, including soil moisture, pH, nutrients and mycorrhizae (Augusto, Dupouey, and Ranger, 2003; Messenger, 1980; Ranger and Claude, 1992) – all of which could affect the phenology of understory plants. Wolf, Zavaleta and Selmants (2017) showed that biotic interactions can affect the timing of flowering, with plants flowering earlier after (experimentally manipulated) biodiversity loss.

### Potential consequences of phenological shifts

A phenology that is fine-tuned to environmental conditions is crucial for plants. Plants that fail to track seasonal temperatures or climatic long-term changes are prone to decline in abundance (Willis et al., 2008). On the other hand, Scheepens and Stöcklin (2013) showed that earlier flowering as a response to climatic changes can also be maladaptive and lead to a fitness decline due to a more rapid development and therefore lower flower numbers. Phenological shifts can alter reproduction and survival, leading to demographic changes (Miller-Rushing et al., 2010), and potentially favouring exotic species (Abu-Asab et al., 2001). Furthermore, a review by Elzinga et al. (2007) argues that that biotic interaction with mutualists and antagonists, e.g. pollinators or pollinator-transmitted fungi, can change plant phenological patterns. It is likely that the biotic and abiotic drivers that determine phenology vary between interacting groups of organisms (or species) such as plants, insects or vertebrates (Parmesan and Yohe, 2003; Voigt et al., 2003). Phenological shifts can alter species interactions and thereby influence the potential for persistence and coexistence of competing species and change biodiversity patterns in natural systems (Rudolf 2019). Asynchronous changes changes could potentially lead to mismatches in phenology (Kharouba et al., 2018; McKinney et al., 2012; Stenseth & Mysterud, 2002; Visser & Both, 2005; Visser, Both, & Lambrechts, 2004), which could exacerbate the effects of climate change on organisms. Several studies found that spring warming can cause plants to flower earlier (Cleland et al., 2007; Parmesan & Yohe, 2003) and create a phenological mismatch between plants and pollinators (Kudo and Ida, 2013; Settele, Bishop, and Potts, 2016), with detrimental effects on plant reproduction (Forrest, 2015) and pollinator fitness (Schenk, Krauss, and Holzschuh, 2018). However, the likeliness of such mismatches is discussed controversially. Renner and Zohner (2018) argue that mismatches due to climate change are most likely in antagonistic interactions, whereas there is only limited evidence of phenological mismatches in mutualistic interactions. A literature review by Kharouba et al. (2018) suggests that a majority (57%) of interacting species changed their phenologies fairly synchronously whereas 43% showed a trend toward asynchrony. Besides affecting the distribution and fitness of interacting species, changes in plant phenology can also affect ecosystem functions such as productivity and carbon cycling, and they can therefore also effect yields in agriculture, horticulture, viticulture, and forestry (Cleland et al., 2007; Menzel et al., 2006).

### Conclusions

Our study shows that plant phenology is affected by forest management. It thus contributes to the growing evidence that, besides climate change, other drivers of current environmental change, such as land use, influence phenology. Forest management interventions – e.g. planting certain tree species, thinning, selective removal of of target trees or even clearfellings – change many forest characteristics such as crown projection area, spatial dispersion of trees and the structural complexity of a forest. Thus, forest management alters forest structure, and thereby changes the microclimatic conditions of a forest stand, its light conditions as well as most likely other environmental factors that impact flowering phenology of understory herbs. These phenology changes in turn can have wide-ranging implications for forest ecosystems and their long-term composition, stability and evolution.

## Acknowledgements

We are grateful to the local management teams in the Schwäbische Alb and Hainich for their assistance in the field. We thank the managers of the three Exploratories, Kirsten Reichel-Jung, Iris Steitz, Sandra Weithmann, Katrin Lorenzen, Juliane Vogt and Miriam Teuscher and all former managers for their work in maintaining the plot and project infrastructure, Christiane Fischer for giving support through the central office, Andreas Ostrowski for managing the central data base, and Markus Fischer, Eduard Linsenmair, Dominik Hessenmöller, Daniel Prati, Ingo Schöning, François Buscot, Ernst-Detlef Schulze, Wolfgang W. Weisser and the late Elisabeth Kalko for their role in setting up the Biodiversity Exploratories project. The work has been (partly) funded by the DFG Priority Program 1374 “Infrastructure-Biodiversity-Exploratories” (DFG project BO 3241/7-1 to OB). Field work permits were issued by the responsible state environmental offices of Baden-Württemberg, Thüringen, and Brandenburg. The authors declare no conflict of interest.

## Author contributions

OB, JFS and AB designed the study. FMW, SB and MS collected the phenology data, and CA and PS contributed the forest management data. FMW compiled and analysed all data with input from OB and JFS. FMW wrote the manuscript with all coauthors contributing to revisions.

## SUPPORTING INFORMATION

**Table S1**: Studied species and their mean flowering start, peak and end as well as the numbers of plots they were flowering on (*N*).

**Table S2**: Relationships between microclimate and the peak flowering of different plant species, listing standardized regression coefficients derived from linear regressions of flowering peak against the different microclimate variables.

**Figure S1**: Regression of flowering peak against mean spring relative humidity.

**Figure S2**: Pearson’s correlations among all forest variables, spring temperature (Ta) and relative humidity (rH) and the respective scatterplots and histograms.

**Table S3**: Relationships between microclimate and the peak flowering of different plant species, listing R^2^-values derived from linear regressions of flowering peak against the different microclimatic variables.

**Table S4**: Relationships between microclimate and the peak flowering of different plant species, listing regression coefficients derived from linear regressions of flowering peak against the different microclimatic variables.

**Table S5**: Relationships between forest characteristics and the peak flowering of different plant species, listing R^2^-values derived from linear regressions of flowering peak against the different forest trait variables.

**Table S6**: Relationships between forest characteristics and the peak flowering of different plant species. The values are regression coefficients derived from linear regressions of flowering peak against the different microclimate variables, with significant values in bold.

**Table S7**: Mean values of estimated management intensity, structural characteristics and microclimatic conditions for the different forest management types.

**Figure S3**: Relationship between crown projection area and spring temperature over 24 hours, during the day and during the night.

**Table S8**: Overview of all explanatory variables that were used within linear regressions and the structural equation model (SEM).

## References

Abu-Asab, M. S., Peterson, P. M., Shetler, S. G., & Orli, S. S. (2001). Earlier plant flowering in spring as a response to global warming in the Washington, DC, area. Biodiversity & Conservation, 10(4), 597–612.

Altermatt, F. (2012). Temperature-related shifts in butterfly phenology depend on the habitat. Global Change Biology, 18(8), 2429–2438.

Anderson, J. T., Inouye, D. W., McKinney, A. M., Colautti, R. I., & Mitchell-Olds, T. (2012). Phenotypic plasticity and adaptive evolution contribute to advancing flowering phenology in response to climate change. Proceedings of the Royal Society B: Biological Sciences, 279(1743), 3843–3852.

Augusto, L., Dupouey, J.-L., & Ranger, J. (2003). Effects of tree species on understory vegetation and environmental conditions in temperate forests. Annals of Forest Science, 60(8), 823–831.

Baker, T. P., Jordan, G. J., Steel, E. A., Fountain-Jones, N. M., Wardlaw, T. J., & Baker, S. C. (2014). Microclimate through space and time: Microclimatic variation at the edge of regeneration forests over daily, yearly and decadal time scales. Forest Ecology and Management, 334, 174–184.

Bellard, C., Bertelsmeier, C., Leadley, P., Thuiller, W., & Courchamp, F. (2012). Impacts of climate change on the future of biodiversity. Ecology Letters, 15(4), 365–377.

Cerdeira Morellato, L. P., Alberton, B., Alvarado, S. T., Borges, B., Buisson, E., Camargo, M. G. G., … Peres, C. A. (2016). Linking plant phenology to conservation biology. Biological Conservation, 195, 60–72.

Chen, J., Saunders, S. C., Crow, T. R., Naiman, R. J., Brosofske, K. D., Mroz, G. D., … Franklin, J. F. (1999). Microclimate in forest ecosystem and landscape ecology: Variations in local climate can be used to monitor and compare the effects of different management regimes. Bioscience, 49(4), 288–297.

Chmielewski, F.-M., Müller, A., & Bruns, E. (2004). Climate changes and trends in phenology of fruit trees and field crops in Germany, 1961–2000. Agricultural and Forest Meteorology, 121(1), 69–78.

Chuine, I. (2010). Why does phenology drive species distribution? Philosophical Transactions of the Royal Society of London. Series B, Biological Sciences, 365(1555), 3149–3160.

Chuine, I., & Regniere, J. (2017). Process-Based Models of Phenology for Plants and Animals. In D. J. Futuyma (Ed.), Annual Review of Ecology, Evolution, and Systematics, 48, 159–182.

Clark, J. S., Salk, C., Melillo, J., & Mohan, J. (2014). Tree phenology responses to winter chilling, spring warming, at north and south range limits. Functional Ecology, 28(6), 1344–1355.

Cleland, E. E., Chuine, I., Menzel, A., Mooney, H. A., & Schwartz, M. D. (2007). Shifting plant phenology in response to global change. Trends in Ecology & Evolution, 22(7), 357–365.

Cook, B. I., Wolkovich, E. M., & Parmesan, C. (2012). Divergent responses to spring and winter warming drive community level flowering trends. Proceedings of the National Academy of Sciences of the United States of America, 109(23), 9000–9005.

Dahlgren, J. P., Zeipel, H. von, & Ehrlén, J. (2007). Variation in vegetative and flowering phenology in a forest herb caused by environmental heterogeneity. American Journal of Botany, 94(9), 1570–1576.

De Frenne, P., Rodríguez-Sánchez, F., Coomes, D. A., Baeten, L., Verstraeten, G., Vellend, M., … Verheyen, K. (2013). Microclimate moderates plant responses to macroclimate warming. Proceedings of the National Academy of Sciences of the United States of America, 110(46), 18561–18565.

del Río, M., Pretzsch, H., Alberdi, I., Bielak, K., Bravo, F., Brunner, A., … Bravo-Oviedo, A. (2016). Characterization of the structure, dynamics, and productivity of mixed-species stands: review and perspectives. European Journal of Forest Research, 135(1), 23–49.

Durant, J. M., Hjermann, D. O., Anker-Nilssen, T., Beaugrand, G., Mysterud, A., Pettorelli, N., & Stenseth, N. C. (2005). Timing and abundance as key mechanisms affecting trophic interactions in variable environments. Ecology Letters, 8(9), 952–958.

Elzinga, J. A., Atlan, A., Biere, A., Gigord, L., Weis, A. E., & Bernasconi, G. (2007). Time after time: flowering phenology and biotic interactions. Trends in Ecology & Evolution, 22(8), 432–439.

Enquist, C. A. F., Kellermann, J. L., Gerst, K. L., & Miller-Rushing, A. J. (2014). Phenology research for natural resource management in the United States. International Journal of Biometeorology, 58(4), 579–589.

Fan, Y., Chen, J., Shirkey, G., John, R., Wu, S. R., Park, H., & Shao, C. (2016). Applications of structural equation modeling (SEM) in ecological studies: an updated review. Ecological Processes, 5(1), 19.

Fischer, M., Bossdorf, O., Gockel, S., Hänsel, F., Hemp, A., Hessenmöller, D., … Weisser, W. W. (2010). Implementing large-scale and long-term functional biodiversity-research: The Biodiversity Exploratories. Basic and Applied Ecology, 11(6), 473–485.

Fitter, A. H., & Fitter, R. S. R. (2002). Rapid changes in flowering time in British plants. Science, 296(5573), 1689–1691.

Fletcher, T. D. (2012). QuantPsyc: Quantitative Psychology Tools. R package version 1.5. Retrieved from https://CRAN.R-project.org/package=QuantPsyc

Forrest, J. R. K. (2015). Plant-pollinator interactions and phenological change: what can we learn about climate impacts from experiments and observations? Oikos, 124(1), 4–13.

Franklin, J., Davis, F. W., Ikegami, M., Syphard, A. D., Flint, L. E., Flint, A. L., & Hannah, L. (2013). Modeling plant species distributions under future climates: how fine scale do climate projections need to be? Global Change Biology, 19(2), 473–483.

Franks, S. J., Sim, S., & Weis, A. E. (2007). Rapid evolution of flowering time by an annual plant in response to a climate fluctuation. Proceedings of the National Academy of Sciences of the United States of America, 104(4), 1278–1282.

Geiger, R., Aron, R. H., & Todhunter, P. (2003). The Climate Near the Ground. Rowman & Littlefield.

Gossner, M. M., Schall, P., Ammer, C., Ammer, U., Engel, K., Schubert, H., … Weisser, W. W. (2014). Forest management intensity measures as alternative to stand properties for quantifying effects on biodiversity. Ecosphere, 5(9), 1–111.

Heikinheimo, M., & Lappalainen, H. (1997). Dependence of the flower bud burst of some plant taxa in Finland on effective temperature sum: implications for climate warming. Annales Botanici Fennici, 34(4), 229–243.

Høye, T. T., Post, Eric, Schmidt, N. M., Trøjelsgaard, K., & Forchhammer, M. C. (2013). Shorter flowering seasons and declining abundance of flower visitors in a warmer Arctic. Nature Climate Change, 3, 759.

Hwang, T., Song, C., Vose, J. M., & Band, L. E. (2011). Topography-mediated controls on local vegetation phenology estimated from MODIS vegetation index. Landscape Ecology, 26(4), 541–556.

IPCC. (2014). Climate Change 2014: Synthesis Report. Contribution of Working Groups I, II and III to the Fifth Assessment Report of the Intergovernmental Panel on Climate Change [Core Writing Team, R.K. Pachauri and L.A. Meyer (eds.)]. IPCC, Geneva, Switzerland, (p. 151 pp.). p. 151 pp.

Kharouba, H. M., Ehrlén, J., Gelman, A., Bolmgren, K., Allen, J. M., Travers, S. E., & Wolkovich, E. M. (2018). Global shifts in the phenological synchrony of species interactions over recent decades. Proceedings of the National Academy of Sciences of the United States of America, 115(20), 5211–5216.

Kline, R. B. (2015). Principles and Practice of Structural Equation Modeling, Fourth Edition. Guilford Publications.

Kudo, G., & Ida, T. Y. (2013). Early onset of spring increases the phenological mismatch between plants and pollinators. Ecology, 94(10), 2311–2320.

Lapointe, L. (2001). How phenology influences physiology in deciduous forest spring ephemerals. Physiologia Plantarum, 113(2), 151–157.

Laube, J., Sparks, T. H., Estrella, N., & Menzel, A. (2014). Does humidity trigger tree phenology? Proposal for an air humidity based framework for bud development in spring. The New Phytologist, 202(2), 350–355.

Lefcheck, J. S. (2016). PiecewiseSEM : Piecewise structural equation modelling in R for ecology, evolution, and systematics. Methods in Ecology and Evolution / British Ecological Society, 7(5), 573–579.

Leong, M., Ponisio, L. C., Kremen, C., Thorp, R. W., & Roderick, G. K. (2016). Temporal dynamics influenced by global change: bee community phenology in urban, agricultural, and natural landscapes. Global Change Biology, 22(3), 1046–1053.

Lieth, H. (1974). Phenology and seasonality modeling. Presented at the New York. Retrieved from http://www.worldcat.org/title/phenology-and-seasonality-modeling/oclc/799774

Matthews, E. R., & Mazer, S. J. (2016). Historical changes in flowering phenology are governed by temperature x precipitation interactions in a widespread perennial herb in western North America. The New Phytologist, 210(1), 157–167.

McKinney, A. M., CaraDonna, P. J., Inouye, D. W., Barr, B., Bertelsen, C. D., & Waser, N. M. (2012). Asynchronous changes in phenology of migrating Broad-tailed Hummingbirds and their early-season nectar resources. Ecology, 93(9), 1987–1993.

Memmott, J., Craze, P. G., Waser, N. M., & Price, M. V. (2007). Global warming and the disruption of plant-pollinator interactions. Ecology Letters, 10(8), 710–717.

Menzel, A., Sparks, T., Estrella, N., Koch, E., & Zust, A. (2006). European phenological response to climate change matches the warming pattern. Global Change Biology, 12(10), 1969.

Messenger, A. S. (1980). Spruce plantation effects on soil moisture and chemical element distribution. Forest Ecology and Management, 3, 113–125.

Miller-Rushing, A. J., Høye, T. T., Inouye, D. W., & Post, E. (2010). The effects of phenological mismatches on demography. Philosophical Transactions of the Royal Society of London. Series B, Biological Sciences, 365(1555), 3177–3186.

Nihlgard, B. (1969). The Microclimate in a beech and a spruce forest - a comparative study from Kongalund, Scania, Sweden,. Bot. Notiser, 5(122), 333–352.

Pacifici, M., Foden, W. B., Visconti, P., Watson, J. E. M., Butchart, S. H. M., Kovacs, K. M., … Rondinini, C. (2015). Assessing species vulnerability to climate change. Nature Climate Change, 5, 215.

Parmesan, C., & Yohe, G. (2003). A globally coherent fingerprint of climate change impacts across natural systems. Nature, 421(6918), 37–42.

Pau, S., Wolkovich, E. M., Cook, B. I., Davies, T. J., Kraft, N. J. B., Bolmgren, K., … Cleland, E. E. (2011). Predicting phenology by integrating ecology, evolution and climate science. Global Change Biology, 17(12), 3633–3643.

Penttinen, A., Stoyan, D., & Henttonen, H. M. (1992). Marked Point Processes in Forest Statistics. Forest Science, 38(4), 806–824.

Ranger, J., & Claude, N. Y. S. (1992). Effects of Spruce Plantation (Picea Abies Karst.) On the Soil Function of a Previous Broadleaved Ecosystem: Analytical and Experimental Investigations. In A. Teller, P. Mathy, & J. N. R. Jeffers (Eds.), Responses of Forest Ecosystems to Environmental Changes (pp. 784–785). Dordrecht: Springer Netherlands.

R Core Team. (2018). R: A language and environment for statistical computing. Vienna: R Foundation for Statistical Computing (Version R version 3.5.0 (2018-04-23)). Retrieved from https://www.R-project.org

Reilly, J., Baethgen, W., Chege, F. E., van de Geijn, S. C., Erda, L., Iglesias, A., … Howden, M. (1996). Agriculture in a changing climate: impacts and adaptation. In Climate change 1995; impacts, adaptations and mitigation of climate change: scientific-technical analyses (pp. 427–467). Cambridge, UK: Cambridge University Press.

Renner, S. S., & Zohner, C. M. (2018). Climate Change and phenological mismatch in trophic interactions among plants, insects, and vertebrates. Annual Review of Ecology, Evolution, and Systematics, 49(1), 165–182.

Richardson, A. D., Anderson, R. S., Arain, M. A., Barr, A. G., Bohrer, G., Chen, G., … Xue, Y. (2012). Terrestrial biosphere models need better representation of vegetation phenology: results from the North American Carbon Program Site Synthesis. Global Change Biology, 18(2), 566–584.

Richardson, A. D., Black, T. A., Ciais, P., Delbart, N., Friedl, M. A., Gobron, N., … Varlagin, A. (2010). Influence of spring and autumn phenological transitions on forest ecosystem productivity. Philosophical Transactions of the Royal Society of London. Series B, Biological Sciences, 365(1555), 3227–3246.

Richardson, A. D., Keenan, T. F., Migliavacca, M., Ryu, Y., Sonnentag, O., & Toomey, M. (2013). Climate change, phenology, and phenological control of vegetation feedbacks to the climate system. Agricultural and Forest Meteorology, 169, 156–173.

Roberts, A. M. I., Tansey, C., Smithers, R. J., & Phillimore, A. B. (2015). Predicting a change in the order of spring phenology in temperate forests. Global Change Biology, 21(7), 2603–2611.

Rudolf, V. H. W. (2019). The role of seasonal timing and phenological shifts for species coexistence. Ecology Letters, 22(8), 1324–1338.

Ryu, E. (2011). Effects of skewness and kurtosis on normal-theory based maximum likelihood test statistic in multilevel structural equation modeling. Behavior Research Methods, 43(4), 1066–1074.

Schall, P., & Ammer, C. (2013). How to quantify forest management intensity in Central European forests. European Journal of Forest Research, 132(2), 379–396.

Schall, P., Schulze, E.-D., Fischer, M., Ayasse, M., & Ammer, C. (2018). Relations between forest management, stand structure and productivity across different types of Central European forests. Basic and Applied Ecology.

Scheepens, J. F., & Stöcklin, J. (2013). Flowering phenology and reproductive fitness along a mountain slope: maladaptive responses to transplantation to a warmer climate in Campanula thyrsoides. Oecologia, 171(3), 679–691.

Schenk, M., Krauss, J., & Holzschuh, A. (2018). Desynchronizations in bee-plant interactions cause severe fitness losses in solitary bees. The Journal of Animal Ecology, 87(1), 139–149.

Schleip, C., Rais, A., & Menzel, A. (2009). Bayesian analysis of temperature sensitivity of plant phenology in Germany. Agricultural and Forest Meteorology, 149(10), 1699–1708.

Schulze, E. D., Aas, G., Grimm, G. W., Gossner, M. M., Walentowski, H., Ammer, C., … von Gadow, K. (2016). A review on plant diversity and forest management of European beech forests. European Journal of Forest Research, 135(1), 51–67.

Schwartz, M. D., Ahas, R., & Aasa, A. (2006). Onset of spring starting earlier across the Northern Hemisphere. Global Change Biology, 12(2), 343–351.

Schweiger, O., Heikkinen, R. K., Harpke, A., Hickler, T., Klotz, S., Kudrna, O., … Settele, J. (2012). Increasing range mismatching of interacting species under global change is related to their ecological characteristics. Global Ecology and Biogeography: A Journal of Macroecology, 21(1), 88–99.

Schweiger, O., Settele, J., Kudrna, O., Klotz, S., & Kühn, I. (2008). Climate change can cause spatial mismatch of trophically interacting species. Ecology, 89(12), 3472–3479.

Settele, J., Bishop, J., & Potts, S. G. (2016). Climate change impacts on pollination. Nature Plants, 2(7), 16092.

Shipley, B. (2009). Confirmatory path analysis in a generalized multilevel context. Ecology, 90(2), 363–368.

Spiecker, H. (2003). Silvicultural management in maintaining biodiversity and resistance of forests in Europe—temperate zone. Journal of Environmental Management, 67(1), 55–65.

Stenseth, N. C., & Mysterud, A. (2002). Climate, changing phenology, and other life history traits: nonlinearity and match-mismatch to the environment. Proceedings of the National Academy of Sciences of the United States of America, 99(21), 13379–13381.

Tang, J., Körner, C., Muraoka, H., Piao, S., Shen, M., Thackeray, S. J., & Yang, X. (2016). Emerging opportunities and challenges in phenology: a review. Ecosphere, 7(8). https://doi.org/10.1002/ecs2.1436

Tinya, F., Márialigeti, S., Király, I., Németh, B., & Ódor, P. (2009). The effect of light conditions on herbs, bryophytes and seedlings of temperate mixed forests in Orség, Western Hungary. Plant Ecology, 204(1), 69.

Visser, M. E., & Both, C. (2005). Shifts in phenology due to global climate change: the need for a yardstick. Proceedings of the Royal Society B: Biological Sciences 272(1581), 2561–2569.

Visser, M. E., Both, C., & Lambrechts, M. M. (2004). Global climate change leads to mistimed avian reproduction. In Advances in Ecological Research (Vol. 35, pp. 89–110). Academic Press.

Voigt, W., Perner, J., Davis, A. J., Eggers, T., Schumacher, J., Bährmann, R., … Sander, F. W. (2003). Trophic levels are differentially sensitive to climate. Ecology, 84(9), 2444–2453.

Völler, E., Auge, H., Bossdorf, O., & Prati, D. (2013). Land use causes genetic differentiation of life-history traits in Bromus hordeaceus. Global Change Biology, 19(3), 892–899.

Völler, E., Bossdorf, O., Prati, D., & Auge, H. (2017). Evolutionary responses to land use in eight common grassland plants. The Journal of Ecology, 105(5), 1290–1297.

von Arx, G., Graf Pannatier, E., Thimonier, A., & Rebetez, M. (2013). Microclimate in forests with varying leaf area index and soil moisture: potential implications for seedling establishment in a changing climate. The Journal of Ecology, 101(5), 1201–1213.

Walther, G.-R. (2010). Community and ecosystem responses to recent climate change. Philosophical Transactions of the Royal Society of London. Series B, Biological Sciences, 365(1549), 2019–2024.

Walther, G.-R., Post, E., Convey, P., Menzel, A., Parmesan, C., Beebee, T. J. C., … Bairlein, F. (2002). Ecological responses to recent climate change. Nature, 416(6879), 389–395.

Wang, L., Tian, F., Wang, Y., Wu, Z., Schurgers, G., & Fensholt, R. (2018). Acceleration of global vegetation greenup from combined effects of climate change and human land management. Global Change Biology, 24(11), 5484–5499.

Ward, S. E., Schulze, M., & Roy, B. (2018). A long-term perspective on microclimate and spring plant phenology in the Western Cascades. Ecosphere, 9(10), e02451.

Willis, C. G., Ruhfel, B., Primack, R. B., Miller-Rushing, A. J., & Davis, C. C. (2008). Phylogenetic patterns of species loss in Thoreau’s woods are driven by climate change. Proceedings of the National Academy of Sciences of the United States of America, 105(44), 17029–17033.

Wolf, A. A., Zavaleta, E. S., & Selmants, P. C. (2017). Flowering phenology shifts in response to biodiversity loss. Proceedings of the National Academy of Sciences of the United States of America, 114(13), 3463–3468.

Wolkovich, E. M., Cook, B. I., Allen, J. M., Crimmins, T. M., Betancourt, J. L., Travers, S. E., … Cleland, E. E. (2012). Warming experiments underpredict plant phenological responses to climate change. Nature, 485(7399), 494–497.

Zenner, E. K. (1998). A new index for describing the structural complexity of forests (Doctoral, Oregon State University). Retrieved from http://ir.library.oregonstate.edu/downloads/bc386n18x

Zenner, E. K., & Hibbs, D. E. (2000). A new method for modeling the heterogeneity of forest structure. Forest Ecology and Management, 129(1), 75–87.

Zhang, X., Liu, L., & Henebry, G. M. (2019). Impacts of land cover and land use change on long-term trend of land surface phenology: a case study in agricultural ecosystems. Environmental Research Letters, 14(4).

Zipf, L., & Primack, R. B. (2017). Humidity does not appear to trigger leaf out in woody plants. International Journal of Biometeorology, 61(12), 2213–2216.

