## Supplemental Figure 1 for "Effects of forest management on the phenology of early-flowering understory herbs"

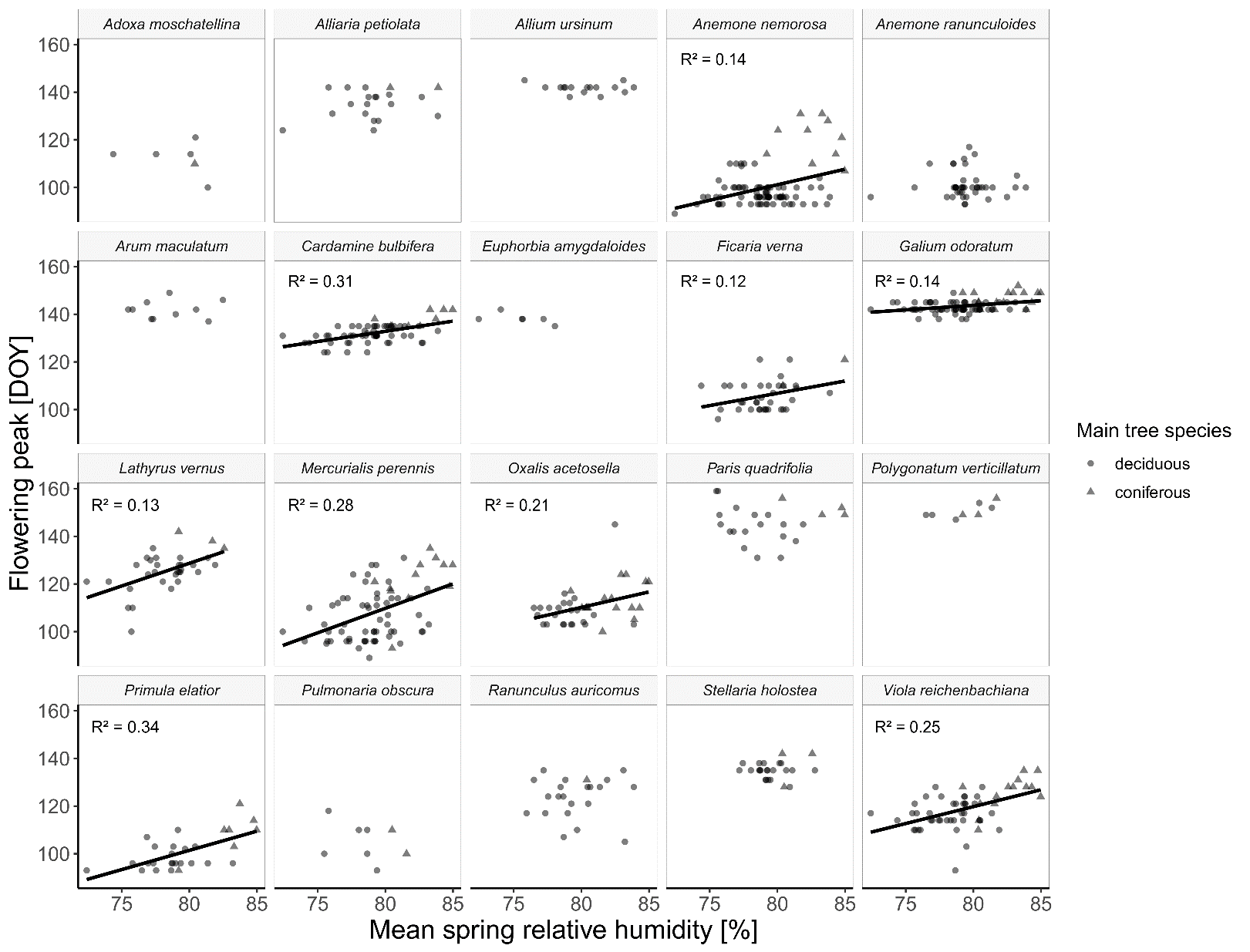

**Figure S1**: Regression of flowering peak against mean spring relative humidity. Each point represents a forest plot, and the shape of each point indicates whether the main tree species is deciduous (circle) or coniferous (triangle). For significant regressions, the regression lines are plotted. All regression coefficients are listed in Table S2.
