## Supplemental Figure 2 for "Effects of forest management on the phenology of early-flowering understory herbs"

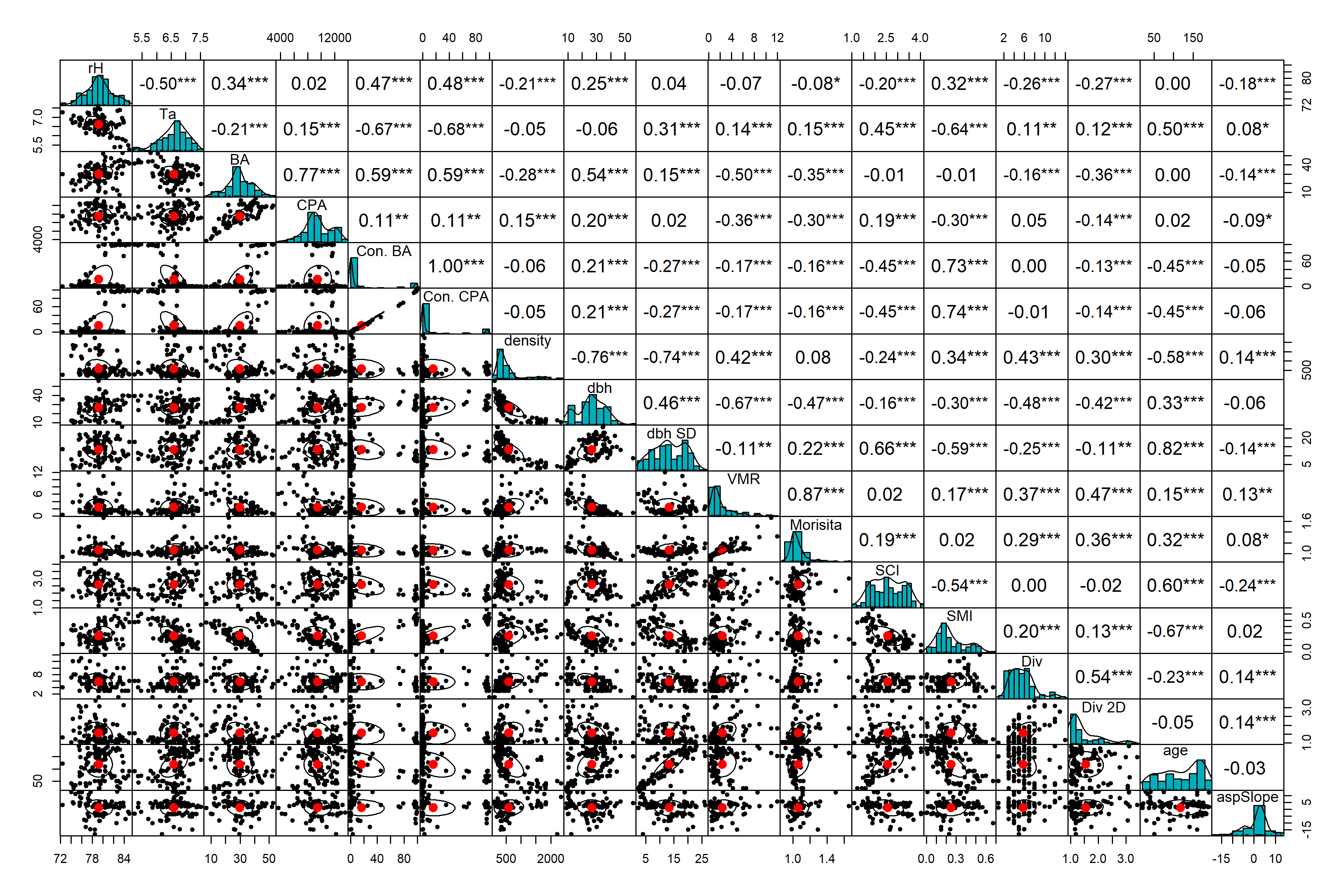

**Figure S2**: Pearson’s correlations among all forest variables, spring temperature (Ta) and relative humidity (rH) and the respective scatterplots and histograms. See Table S8 for details on the variables. Significance levels: * *P* < 0.05, ** *P* < 0.01, *** *P* < 0.001.
