## Supplemental Table 1 for "Effects of forest management on the phenology of early-flowering understory herbs"

**Table S1**: Studied species and their mean flowering start, peak and end as well as the numbers of plots they were flowering on (*N*).

| **Species** | **Flowering start** | | **Flowering peak** | | **Flowering end** | |
| --- | --- | --- | --- | --- | --- | --- |
| *Adoxa moschatellina* | April 8 | (7) | April 22 | (6) | May 2 | (6) |
| *Alliaria petiolata* | May 4 | (22) | May 15 | (20) | May 30 | (18) |
| *Allium ursinum* | May 12 | (16) | May 22 | (16) | June 1 | (16) |
| *Anemone nemorosa* | March 27 | (88) | April 10 | (87) | May 16 | (87) |
| *Anemone ranunculoides* | March 31 | (43) | April 11 | (42) | May 11 | (41) |
| *Arum maculatum* | May 17 | (11) | May 22 | (10) | May 31 | (10) |
| *Cardamine bulbifera* | May 5 | (60) | May 12 | (62) | May 22 | (62) |
| *Euphorbia amygdaloides* | April 20 | (10) | May 18 | (6) | June 12 | (3) |
| *Ficaria verna* | April 7 | (43) | April 16 | (37) | May 3 | (36) |
| *Galium odoratum* | May 12 | (79) | May 24 | (79) | June 12 | (79) |
| *Lathyrus vernus* | April 23 | (35) | May 5 | (32) | May 18 | (31) |
| *Mercurialis perennis* | March 26 | (66) | April 19 | (71) | May 16 | (71) |
| *Oxalis acetosella* | April 8 | (50) | April 21 | (39) | May 13 | (39) |
| *Paris quadrifolia* | May 11 | (24) | May 26 | (22) | June 8 | (9) |
| *Polygonatum verticillatum* | May 26 | (9) | May 31 | (8) | June 5 | (8) |
| *Primula elatior* | March 29 | (25) | April 11 | (28) | May 1 | (29) |
| *Pulmonaria obscura* | April 4 | (9) | April 15 | (8) | May 13 | (8) |
| *Ranunculus auricomus* | April 22 | (22) | May 4 | (22) | May 17 | (22) |
| *Stellaria holostea* | April 28 | (24) | May 15 | (24) | May 31 | (22) |
| *Viola reichenbachiana* | April 11 | (58) | April 29 | (55) | May 16 | (54) |
