## Supplemental Table 3 for "Effects of forest management on the phenology of early-flowering understory herbs"

**Table S3**: Relationships between microclimate and the peak flowering of different plant species. The values are R^2^-values derived from linear regressions of flowering peak against the different microclimatic variables. The winter period encompasses October 2016 – January 2017 and the spring period February – May 2017. Ice days = the number of days with a maximum temperature < 0°C, cold days = the number of days with a temperature minimum < 0°C, cold sum = the sum of days with a mean day temperature < 0 °C, cool days = the number of days with a temperature maximum < 10°C, Ta = air temperature in °C, rH = relative humidity, Ts = soil temperature, SM = soil moisture, GD = days with temperatures between 10°C to 30°C, growth sum = the sum of mean day temperature > 5°C, warm sum = sum of day temperatures with a mean > 10 °C.

|  | **Winter** | | | | | | | | | | **Summer** | | | | | | | |
| --- | --- | --- | --- | --- | --- | --- | --- | --- | --- | --- | --- | --- | --- | --- | --- | --- | --- | --- |
|  | **Ice days** | **Cold days** | **Cold sum** | **Cool days** | **Ta**  **10 cm** | **Ta**  **200 cm** | **Ts** | **rH** | **SM** |  | **GD** | **growth sum** | **warm sum** | **Ta**  **10 cm** | **Ta**  **200 cm** | **Ts** | **rH** | **SM** |
| *Alliaria petiolata* | 0.10 | 0.20 | 0.25 | 0.00 | 0.08 | 0.32 | 0.02 | 0.03 | 0.11 |  | 0.07 | 0.38 | 0.29 | 0.14 | 0.41 | 0.01 | 0.07 | 0.13 |
| *Allium ursinum* | 0.17 | 0.04 | 0.06 | 0.17 | 0.04 | 0.00 | 0.02 | 0.05 | 0.05 |  | 0.08 | 0.00 | 0.01 | 0.00 | 0.00 | 0.02 | 0.03 | 0.03 |
| *Anemone nemorosa* | 0.10 | 0.28 | 0.35 | 0.00 | 0.14 | 0.36 | 0.02 | 0.01 | 0.01 |  | 0.37 | 0.39 | 0.45 | 0.28 | 0.39 | 0.00 | 0.14 | 0.04 |
| *Anemone ranunculoides* | 0.01 | 0.03 | 0.09 | 0.00 | 0.01 | 0.05 | 0.00 | 0.00 | 0.02 |  | 0.02 | 0.01 | 0.03 | 0.07 | 0.01 | 0.01 | 0.00 | 0.03 |
| *Arum maculatum* | 0.09 | 0.26 | 0.09 | 0.00 | 0.01 | 0.14 | 0.02 | 0.01 | 0.04 |  | 0.14 | 0.14 | 0.26 | 0.04 | 0.09 | 0.00 | 0.00 | 0.00 |
| *Cardamine bulbifera* | 0.10 | 0.14 | 0.10 | 0.17 | 0.20 | 0.18 | 0.09 | 0.13 | 0.04 |  | 0.27 | 0.44 | 0.32 | 0.28 | 0.43 | 0.17 | 0.31 | 0.01 |
| *Ficaria verna* | 0.10 | 0.28 | 0.33 | 0.00 | 0.14 | 0.38 | 0.03 | 0.05 | 0.01 |  | 0.22 | 0.42 | 0.43 | 0.19 | 0.44 | 0.01 | 0.12 | 0.00 |
| *Galium odoratum* | 0.08 | 0.29 | 0.37 | 0.01 | 0.20 | 0.41 | 0.00 | 0.00 | 0.00 |  | 0.35 | 0.53 | 0.52 | 0.43 | 0.52 | 0.02 | 0.13 | 0.02 |
| *Lathyrus vernus* | 0.11 | 0.14 | 0.12 | 0.03 | 0.11 | 0.16 | 0.01 | 0.10 | 0.01 |  | 0.28 | 0.27 | 0.25 | 0.20 | 0.27 | 0.06 | 0.28 | 0.01 |
| *Mercurialis perennis* | 0.03 | 0.31 | 0.37 | 0.00 | 0.24 | 0.38 | 0.04 | 0.05 | 0.00 |  | 0.31 | 0.38 | 0.43 | 0.37 | 0.37 | 0.10 | 0.21 | 0.00 |
| *Oxalis acetosella* | 0.00 | 0.13 | 0.16 | 0.00 | 0.19 | 0.20 | 0.00 | 0.01 | 0.01 |  | 0.11 | 0.24 | 0.23 | 0.27 | 0.24 | 0.03 | 0.14 | 0.01 |
| *Paris quadrifolia* | 0.06 | 0.00 | 0.01 | 0.04 | 0.00 | 0.00 | 0.00 | 0.05 | 0.04 |  | 0.00 | 0.02 | 0.03 | 0.01 | 0.01 | 0.05 | 0.01 | 0.09 |
| *Polygonatum verticillatum* | 0.59 | 0.04 | 0.07 | 0.07 | 0.00 | 0.13 | 0.01 | 0.63 | 0.09 |  | 0.13 | 0.14 | 0.06 | 0.04 | 0.21 | 0.02 | 0.47 | 0.14 |
| *Primula elatior* | 0.29 | 0.20 | 0.15 | 0.12 | 0.01 | 0.18 | 0.01 | 0.25 | 0.00 |  | 0.11 | 0.26 | 0.23 | 0.19 | 0.27 | 0.03 | 0.40 | 0.00 |
| *Pulmonaria obscura* | 0.07 | 0.00 | 0.00 | 0.02 | 0.59 | 0.01 | 0.29 | 0.23 | 0.05 |  | 0.16 | 0.01 | 0.02 | 0.34 | 0.00 | 0.11 | 0.10 | 0.02 |
| *Ranunculus auricomus* | 0.19 | 0.28 | 0.27 | 0.00 | 0.15 | 0.36 | 0.03 | 0.00 | 0.00 |  | 0.26 | 0.44 | 0.29 | 0.23 | 0.46 | 0.00 | 0.00 | 0.00 |
| *Stellaria holostea* | 0.01 | 0.03 | 0.11 | 0.07 | 0.07 | 0.07 | 0.08 | 0.01 | 0.03 |  | 0.14 | 0.02 | 0.04 | 0.02 | 0.02 | 0.06 | 0.01 | 0.01 |
| *Viola reichenbachiana* | 0.24 | 0.16 | 0.16 | 0.12 | 0.11 | 0.20 | 0.02 | 0.11 | 0.01 |  | 0.24 | 0.31 | 0.22 | 0.22 | 0.33 | 0.03 | 0.25 | 0.01 |
| Average over all species | 0.13 | 0.15 | 0.17 | 0.05 | 0.13 | 0.20 | 0.04 | 0.09 | 0.03 |  | 0.18 | 0.24 | 0.23 | 0.18 | 0.25 | 0.04 | 0.15 | 0.03 |
