## Supplemental Table 4 for "Effects of forest management on the phenology of early-flowering understory herbs"

**Table S4**: Relationships between microclimate and the peak flowering of different plant species. The values are regression coefficients derived from linear regressions of flowering peak against the different microclimatic variables, with significant values in bold. The winter period encompasses October 2016 – January 2017 and the spring period February – May 2017. Ice days = the number of days with a maximum temperature < 0°C, cold days = the number of days with a temperature minimum < 0°C, cold sum = the sum of days with a mean day temperature < 0 °C, cool days = the number of days with a temperature maximum < 10°C, Ta = air temperature in °C, rH = relative humidity, Ts = soil temperature, SM = soil moisture, GD = days with temperatures between 10°C to 30°C, growth sum = the sum of mean day temperature > 5°C, warm sum = sum of day temperatures with a mean > 10 °C.

|  | **Winter** | | | | | | | | |  | **Spring** | | | | | | | |
| --- | --- | --- | --- | --- | --- | --- | --- | --- | --- | --- | --- | --- | --- | --- | --- | --- | --- | --- |
|  | **Ice**  **days** | **Cold**  **days** | **Cold**  **sum** | **Cool**  **days** | **Ta**  **10 cm** | **Ta**  **200 cm** | **Ts** | **rH** | **SM** |  | **GD** | **growth**  **sum** | **warm**  **sum** | **Ta 10 cm** | **Ta 200 cm** | **Ts** | **rH** | **SM** |
| *Alliaria petiolata* | 0.49 | 0.47 | **0.11** | 0.05 | -4.70 | **-10.89** | -0.65 | 0.58 | 0.50 |  | -0.70 | **-0.13** | **-0.22** | -3.02 | **-7.29** | -0.86 | 0.32 | 0.49 |
| *Allium ursinum* | -0.19 | 0.14 | 0.04 | -0.23 | -0.05 | 0.34 | 0.65 | -0.14 | 0.08 |  | -0.42 | 0.00 | -0.02 | -1.01 | -0.08 | -0.41 | -0.19 | 0.11 |
| *Anemone nemorosa* | **0.70** | 0.70 | **0.15** | 0.09 | **-8.70** | **-11.45** | -0.44 | 1.31 | -0.30 |  | **-1.20** | **-0.14** | **-0.35** | **-5.81** | **-9.12** | 1.09 | **0.33** | -0.15 |
| *Anemone ranunculoides* | 0.15 | 0.19 | 0.07 | -0.12 | -3.48 | -1.77 | -0.84 | 0.03 | 0.20 |  | -0.22 | -0.02 | -0.09 | -1.04 | -3.08 | 0.13 | -0.02 | 0.13 |
| *Arum maculatum* | -0.30 | 0.40 | 0.05 | 0.04 | -1.99 | -3.17 | 0.07 | 0.11 | -0.01 |  | -0.56 | -0.05 | -0.17 | 0.73 | -3.70 | 0.52 | -0.11 | -0.10 |
| *Cardamine bulbifera* | **0.33** | 0.22 | **0.04** | **0.33** | **-3.96** | **-6.05** | **-2.39** | **0.86** | 0.07 |  | **-0.44** | **-0.08** | **-0.14** | **-2.84** | **-3.08** | **-1.40** | **0.53** | 0.14 |
| *Ficaria verna* | 0.52 | 0.51 | **0.10** | 0.02 | **-5.14** | **-9.52** | -0.75 | 1.03 | 0.08 |  | **-0.76** | **-0.12** | **-0.26** | **-4.05** | **-6.89** | -1.04 | **0.61** | 0.14 |
| *Galium odoratum* | **0.22** | 0.22 | **0.05** | 0.05 | **-3.30** | **-4.41** | -0.52 | 0.38 | -0.07 |  | **-0.35** | **-0.06** | **-0.12** | **-2.07** | **-3.24** | 0.08 | **0.04** | -0.03 |
| *Lathyrus vernus* | 0.72 | 0.48 | 0.09 | 0.29 | -6.83 | **-10.95** | -2.06 | 1.91 | 0.09 |  | **-1.12** | **-0.14** | **-0.28** | **-3.96** | **-6.35** | -0.52 | **0.94** | 0.10 |
| *Mercurialis perennis* | 0.54 | 0.90 | **0.20** | -0.13 | **-13.68** | **-14.81** | -4.85 | 2.06 | 0.06 |  | **-1.36** | **-0.19** | **-0.43** | **-9.37** | **-12.32** | **-2.41** | **0.97** | 0.11 |
| *Oxalis acetosella* | -0.11 | 0.41 | **0.08** | -0.14 | **-6.83** | **-7.27** | -1.76 | 1.30 | 0.11 |  | **-0.55** | **-0.09** | **-0.19** | **-6.24** | **-5.74** | -0.55 | **0.40** | 0.11 |
| *Paris quadrifolia* | -0.49 | 0.00 | 0.02 | -0.28 | 1.45 | -1.71 | 3.07 | -0.19 | -0.41 |  | 0.04 | -0.03 | -0.07 | -0.06 | -0.87 | 0.67 | -0.70 | -0.28 |
| *Polygonatum verticillatum* | **0.93** | 0.10 | 0.03 | 0.35 | -1.34 | -4.01 | 0.64 | **1.09** | -0.17 |  | -0.24 | -0.04 | -0.04 | -0.22 | -2.56 | 0.22 | 1.29 | -0.14 |
| *Primula elatior* | **0.95** | 0.39 | **0.07** | 0.76 | -4.74 | **-6.15** | -1.60 | **1.61** | -0.05 |  | -0.44 | **-0.08** | **-0.16** | **-1.29** | **-4.20** | 0.62 | **1.33** | 0.05 |
| *Pulmonaria obscura* | 0.56 | 0.07 | 0.00 | -0.22 | **-13.95** | -0.10 | -2.69 | -1.24 | -0.13 |  | -1.10 | -0.02 | -0.11 | -12.79 | 3.64 | -4.32 | -1.54 | -0.22 |
| *Ranunculus auricomus* | **1.05** | 0.64 | **0.13** | 0.00 | -8.06 | **-14.92** | -0.62 | 0.24 | 0.06 |  | **-1.12** | **-0.18** | **-0.33** | **-5.68** | **-9.41** | -1.46 | 0.08 | -0.02 |
| *Stellaria holostea* | 0.10 | 0.11 | 0.05 | -0.37 | -1.01 | -1.21 | -0.96 | 0.20 | -0.03 |  | -0.32 | -0.02 | -0.05 | -1.61 | -2.27 | -0.97 | -0.27 | -0.09 |
| *Viola reichenbachiana* | **1.10** | 0.46 | **0.09** | **0.56** | **-6.35** | **-9.49** | -1.61 | **1.42** | -0.10 |  | **-0.84** | **-0.11** | **-0.20** | **-3.46** | **-6.15** | 0.96 | **0.90** | 0.09 |
| Average over all species | 0.40 | 0.36 | 0.08 | 0.06 | -5.15 | -6.53 | -0.96 | 0.70 | 0.00 |  | -0.65 | -0.08 | -0.18 | -3.54 | -4.59 | -0.54 | 0.27 | 0.02 |
