## Supplemental Table 5 for "Effects of forest management on the phenology of early-flowering understory herbs"

**Table S5**: Relationships between forest characteristics and the peak flowering of different plant species. The values are R^2^-values derived from linear regressions of flowering peak against the different forest variables. Age = mean age of the main tree species, AspSlope = the inclination multiplied by 1 for south-, 1 for north-, and 0.5 for east- and west-facing slopes, BA = basal area covered with trees in m^2^ ha^−1^, CPA = crown projection area in m^2^ ha^−^1, Con = Coniferous, dbh = diameter at breast height in cm, dbh SD = the standard deviation of dbh, density = stand density (trees ha^−1^), Div = species richness, Div 2D = inverse Simpson’s index, Morisita = Morisita's index of dispersion, SCI = Zenner’s Structural Complexity Index (SCI) based on tree height, SMI = silvicultural management intensity, VMR= Clapham’s variance mean ratio (VMR).

|  | **Age** | **AspSlope** | **BA** | **Con BA** | **Con CPA** | **CPA** | **Dbh** | **Dbh SD** | **Density** | **Div** | **Div 2D** | **Morisita** | **SCI** | **SMI** | **VMR** |
| --- | --- | --- | --- | --- | --- | --- | --- | --- | --- | --- | --- | --- | --- | --- | --- |
| *Alliaria petiolata* | 0.166 | 0.006 | 0.01 | 0.178 | 0.171 | 0.003 | 0.002 | 0.074 | 0.053 | 0.101 | 0.07 | 0.037 | 0.07 | 0.164 | 0.027 |
| *Allium ursinum* | 0.272 | 0.018 | 0.181 |  |  | 0.149 | 0.009 | 0.086 | 0.036 | 0.057 | 0.297 | 0.019 | 0.109 | 0.066 | 0.023 |
| *Anemone nemorosa* | 0.136 | 0 | 0.208 | 0.668 | 0.669 | 0.007 | 0.025 | 0.046 | 0.002 | 0.018 | 0.029 | 0.017 | 0.113 | 0.288 | 0.017 |
| *Anemone ranunculoides* | 0.012 | 0.169 | 0 | 0.104 | 0.118 | 0.006 | 0.038 | 0.039 | 0.08 | 0.04 | 0.001 | 0.006 | 0.005 | 0.005 | 0.048 |
| *Arum maculatum* | 0.303 | 0.401 | 0.009 | 0.036 | 0.032 | 0.003 | 0.131 | 0.121 | 0.325 | 0.012 | 0.003 | 0.003 | 0.049 | 0.048 | 0.015 |
| *Cardamine bulbifera* | 0.047 | 0.111 | 0.226 | 0.339 | 0.351 | 0.013 | 0.094 | 0 | 0.049 | 0.098 | 0.122 | 0.124 | 0.043 | 0.102 | 0.138 |
| *Ficaria verna* | 0.098 | 0 | 0 | 0.139 | 0.146 | 0.027 | 0 | 0.03 | 0.007 | 0.001 | 0.003 | 0.034 | 0.13 | 0.203 | 0.025 |
| *Galium odoratum* | 0.15 | 0.056 | 0.132 | 0.365 | 0.373 | 0.007 | 0.019 | 0.035 | 0.001 | 0.045 | 0.042 | 0.029 | 0.173 | 0.237 | 0.034 |
| *Lathyrus vernus* | 0.019 | 0.079 | 0.016 | 0.254 | 0.263 | 0.081 | 0.007 | 0.028 | 0.047 | 0.08 | 0.16 | 0.011 | 0.14 | 0.173 | 0.014 |
| *Mercurialis perennis* | 0.151 | 0.051 | 0.035 | 0.276 | 0.283 | 0.007 | 0.004 | 0.073 | 0 | 0.003 | 0.006 | 0.048 | 0.099 | 0.226 | 0.023 |
| *Oxalis acetosella* | 0.024 | 0.013 | 0.002 | 0.078 | 0.082 | 0.058 | 0.001 | 0.006 | 0.001 | 0.042 | 0.018 | 0.016 | 0.033 | 0.09 | 0.013 |
| *Paris quadrifolia* | 0.018 | 0.24 | 0.036 | 0.199 | 0.183 | 0.02 | 0.058 | 0.004 | 0.015 | 0.058 | 0.002 | 0 | 0.16 | 0.17 | 0 |
| *Polygonatum verticillatum* | 0.046 | 0.094 | 0.01 | 0.023 | 0.027 | 0.001 | 0.005 | 0.157 | 0.007 | 0.2 | 0.039 | 0.002 | 0.112 | 0.01 | 0.002 |
| *Primula elatior* | 0.066 | 0.009 | 0.208 | 0.392 | 0.402 | 0.002 | 0.056 | 0.01 | 0.008 | 0.062 | 0.036 | 0.018 | 0.043 | 0.32 | 0.012 |
| *Pulmonaria obscura* | 0.199 | 0.001 | 0.051 | 0.003 | 0.006 | 0.037 | 0.124 | 0.214 | 0.138 | 0.071 | 0.005 | 0.079 | 0.631 | 0.21 | 0.128 |
| *Ranunculus auricomus* | 0.086 | 0.058 | 0.061 | 0.053 | 0.052 | 0.13 | 0.012 | 0.038 | 0.01 | 0.05 | 0.056 | 0.061 | 0.101 | 0.067 | 0.021 |
| *Stellaria holostea* | 0.142 | 0.178 | 0.004 | 0.102 | 0.122 | 0.002 | 0.054 | 0.177 | 0.045 | 0.074 | 0.002 | 0.007 | 0.037 | 0.077 | 0.027 |
| *Viola reichenbachiana* | 0.059 | 0.079 | 0.199 | 0.263 | 0.27 | 0.006 | 0.098 | 0 | 0.059 | 0.128 | 0.166 | 0.057 | 0.04 | 0.099 | 0.11 |
| Average over all species | 0.11 | 0.09 | 0.08 | 0.20 | 0.21 | 0.03 | 0.04 | 0.06 | 0.05 | 0.06 | 0.06 | 0.03 | 0.12 | 0.14 | 0.04 |
