## Supplemental Table 6 for "Effects of forest management on the phenology of early-flowering understory herbs"

**Table S6**: Relationships between forest characteristics and the peak flowering of different plant species. The values are regression coefficients derived from linear regressions of flowering peak against the different forest variables, with significant values in bold. Significant values are written in bold. Age = mean age of the main tree species, AspSlope = the inclination multiplied by 1 for south-, 1 for north-, and 0.5 for east- and west-facing slopes, BA = basal area covered with trees in m^2^ ha^−1^, CPA = crown projection area in m^2^ ha^−^1, Con = Coniferous, dbh = diameter at breast height in cm, dbh SD = the standard deviation of dbh, density = stand density (trees ha^−1^), Div = species richness, Div 2D = inverse Simpson’s index, Morisita = Morisita's index of dispersion, SCI = Zenner’s Structural Complexity Index (SCI) based on tree height, SMI = silvicultural management intensity, VMR= Clapham’s variance mean ratio (VMR).

|  | **Age** | **AspSlope** | **BA** | **Con BA** | **Con CPA** | **CPA** | **Dbh** | **Dbh SD** | **Density** | **Div** | **Div 2D** | **Morisita** | **SCI** | **SMI** | **VMR** |
| --- | --- | --- | --- | --- | --- | --- | --- | --- | --- | --- | --- | --- | --- | --- | --- |
| *Alliaria petiolata* | -0.05 | -0.06 | 0.09 | 0.10 | 0.10 | 0.00 | -0.02 | -0.27 | 0.00 | 0.63 | 2.03 | -13.82 | -2.13 | 20.71 | -0.75 |
| *Allium ursinum* | -0.02 | 0.09 | -0.10 |  |  | 0.00 | -0.02 | -0.11 | 0.00 | 0.32 | **2.20** | -5.15 | -1.08 | 4.35 | 0.22 |
| *Anemone nemorosa* | -0.06 | 0.01 | **0.45** | **0.23** | **0.24** | 0.00 | 0.14 | **-0.33** | 0.00 | -0.55 | -2.63 | -10.00 | **-4.57** | **31.56** | -0.49 |
| *Anemone ranunculoides* | -0.01 | -0.44 | 0.00 | **0.78** | **1.17** | 0.00 | -0.11 | -0.19 | 0.00 | 0.64 | 0.30 | 3.37 | -0.51 | 3.18 | 0.51 |
| *Arum maculatum* | -0.04 | -0.28 | -0.04 | 0.17 | 0.17 | 0.00 | -0.15 | -0.26 | 0.00 | 0.16 | 0.26 | 0.95 | 1.34 | 6.52 | -0.09 |
| *Cardamine bulbifera* | -0.02 | -0.24 | **0.22** | **0.07** | **0.08** | 0.00 | **0.12** | -0.01 | 0.00 | **-0.51** | **-2.32** | **-10.94** | -1.21 | **9.01** | **-0.64** |
| *Ficaria verna* | -0.05 | -0.03 | 0.01 | **0.12** | **0.13** | 0.00 | 0.01 | -0.22 | 0.00 | -0.08 | -0.65 | -19.75 | **-3.91** | **26.42** | -0.79 |
| *Galium odoratum* | -0.02 | -0.10 | **0.10** | **0.05** | **0.05** | 0.00 | 0.04 | -0.09 | 0.00 | -0.23 | -0.96 | -3.95 | **-1.81** | **9.05** | -0.23 |
| *Lathyrus vernus* | -0.02 | -0.36 | 0.15 | **0.15** | **0.15** | 0.00 | 0.07 | -0.25 | 0.00 | -0.91 | **-4.79** | -5.64 | -3.73 | **22.42** | 0.30 |
| *Mercurialis perennis* | -0.09 | -0.42 | 0.24 | **0.18** | **0.18** | 0.00 | 0.08 | **-0.55** | 0.00 | -0.26 | -1.49 | -21.58 | **-5.53** | **34.55** | -0.85 |
| *Oxalis acetosella* | -0.03 | 0.21 | 0.05 | 0.05 | 0.05 | 0.00 | 0.03 | -0.13 | 0.00 | -0.70 | -2.55 | -9.00 | -2.14 | 13.74 | -0.50 |
| *Paris quadrifolia* | -0.02 | 0.56 | 0.17 | **0.09** | **0.09** | 0.00 | 0.19 | -0.09 | 0.00 | 0.63 | 0.50 | 1.20 | -4.88 | 22.96 | 0.06 |
| *Polygonatum verticillatum* | -0.02 | 0.23 | 0.05 | 0.01 | 0.01 | 0.00 | -0.04 | -0.40 | 0.00 | -0.38 | -1.98 | 4.05 | -1.37 | 1.59 | -0.14 |
| *Primula elatior* | -0.04 | -0.15 | **0.41** | **0.11** | **0.11** | 0.00 | 0.24 | -0.17 | 0.00 | -0.83 | -2.51 | -7.27 | -2.20 | **22.75** | -0.37 |
| *Pulmonaria obscura* | 0.07 | 0.06 | 0.16 | -0.01 | -0.02 | 0.00 | 0.25 | 0.65 | -0.01 | -1.17 | 0.60 | -13.22 | **10.46** | -23.29 | -1.03 |
| *Ranunculus auricomus* | -0.05 | 0.46 | -0.20 | 0.11 | 0.13 | 0.00 | 0.09 | -0.29 | 0.00 | -0.73 | -3.32 | -23.54 | -3.61 | 13.96 | -0.50 |
| *Stellaria holostea* | -0.03 | -0.40 | 0.03 | 0.04 | 0.04 | 0.00 | -0.09 | **-0.29** | 0.00 | 0.37 | 0.33 | 2.15 | -0.97 | 6.37 | 0.25 |
| *Viola reichenbachiana* | -0.04 | -0.36 | **0.36** | **0.10** | **0.10** | 0.00 | **0.23** | 0.02 | 0.00 | **-0.96** | **-4.65** | -17.68 | -2.30 | **16.09** | **-1.24** |
| Average over all species | -0.03 | -0.07 | 0.12 | 0.14 | 0.16 | 0.00 | 0.06 | -0.17 | 0.00 | -0.25 | -1.20 | -8.32 | -1.68 | 13.44 | -0.35 |
