## Supplemental Table 7 for "Effects of forest management on the phenology of early-flowering understory herbs"

**Table S7**: Mean values of estimated management intensity, structural characteristics and microclimatic conditions for the different forest management types. Higher colour saturation indicates higher values. AC = age class forest consisting mainly of trees of the same age, Age = mean age of the main tree species, BA = basal area covered with trees in m^2^ ha^−1^, CPA = crown projection area in m^2^ ha^−^1, Con = Coniferous, dbh = diameter at breast height in cm, dbh SD = the standard deviation of dbh, density = stand density (trees ha^−1^), Div = species richness, Div 2D = inverse Simpson’s index, Morisita = Morisita's index of dispersion, SCI = Zenner’s Structural Complexity Index (SCI) based on tree height, SMI = silvicultural management intensity, VMR= Clapham’s variance mean ratio (VMR). Ta = air temperature in °C and rH = relative humidity, both measured during the spring period February – May 2017 at 200 cm height.

| **Forest management type** | **N** |  | **SMI** | **Age** | **BA** | **Con BA** | **Con CPA** | **CPA** | **Density** | **Dbh** | **Dbh SD** | **Div** | **Div 2D** | **Morisita** | **SCI** | **VMR** |  | **Ta** | **rH** |
| --- | --- | --- | --- | --- | --- | --- | --- | --- | --- | --- | --- | --- | --- | --- | --- | --- | --- | --- | --- |
| Beech unmanaged mature timber | 18 |  | 0.10 | 173.93 | 35.68 | 2.12 | 1.67 | 11607.10 | 447.15 | 27.03 | 18.39 | 6.38 | 1.62 | 1.12 | 3.06 | 3.14 |  | 6.99 | 78.93 |
| Beech AC mature timber | 11 |  | 0.16 | 138.39 | 28.84 | 0.33 | 0.23 | 9118.52 | 236.87 | 37.23 | 15.33 | 4.46 | 1.47 | 1.01 | 2.17 | 1.30 |  | 6.75 | 78.70 |
| Beech AC immature timber | 8 |  | 0.17 | 101.39 | 30.58 | 3.12 | 2.51 | 9732.36 | 349.27 | 31.61 | 12.41 | 4.65 | 1.20 | 0.99 | 2.59 | 0.82 |  | 6.57 | 78.54 |
| Beech selection system | 13 |  | 0.17 | 171.32 | 27.46 | 0.00 | 0.00 | 8710.53 | 300.76 | 29.18 | 19.47 | 4.59 | 1.17 | 1.11 | 3.17 | 2.59 |  | 6.80 | 79.90 |
| Beech AC mixed immature timber | 7 |  | 0.23 | 102.78 | 29.39 | 22.07 | 17.65 | 8988.78 | 431.62 | 27.33 | 13.03 | 5.41 | 1.90 | 1.01 | 2.85 | 1.12 |  | 6.52 | 77.92 |
| Beech AC pole wood | 10 |  | 0.29 | 69.42 | 26.82 | 0.93 | 0.77 | 11124.93 | 1679.36 | 13.22 | 5.60 | 8.21 | 1.86 | 1.05 | 2.35 | 4.56 |  | 6.66 | 77.33 |
| Beech AC thicket with shelterwood | 3 |  | 0.34 | 158.25 | 18.11 | 0.54 | 0.33 | 6470.21 | 534.42 | 16.20 | 15.35 | 4.83 | 2.12 | 1.15 | 2.74 | 3.93 |  | 6.50 | 79.24 |
| Beech AC thicket | 13 |  | 0.39 | 71.76 | 14.91 | 2.82 | 2.60 | 6803.31 | 1197.95 | 11.07 | 7.04 | 7.89 | 2.15 | 1.16 | 1.96 | 6.94 |  | 6.59 | 78.66 |
| Spruce AC mature timber | 5 |  | 0.48 | 84.26 | 46.47 | 94.43 | 91.39 | 10470.38 | 326.12 | 40.49 | 13.69 | 5.71 | 1.39 | 0.99 | 2.06 | 0.78 |  | 5.69 | 83.19 |
| Spruce AC immature timber | 10 |  | 0.54 | 56.01 | 40.76 | 95.90 | 93.95 | 10082.35 | 649.43 | 27.41 | 8.23 | 5.94 | 1.27 | 1.04 | 1.81 | 2.09 |  | 5.94 | 82.01 |
