## Supplemental Table 8 for "Effects of forest management on the phenology of early-flowering understory herbs"

**Table S8**: Overview of all explanatory variables that were used within linear regressions and the structural equation model (SEM).

| **Variable** | **Explanation** | **Unit** | **In SEM** |
| --- | --- | --- | --- |
| **Forest variables** |  |  |  |
| Age | Mean age of the main tree species | years |  |
| Basal area | Basal area covered with trees | m^2^ ha^-1^ |  |
| Clapham’s variance mean ratio | Horizontal dispersion, <1: regular, >1: clumping, 1: random; 20 m × 20 m raster cells |  | x |
| Crown projection area | Cumulative crown projection area of trees | m^2^ ha^-1^ | x |
| Coniferous basal area | Percentage of conifers based on basal area | m^2^ ha^-1^ | x |
| Coniferous crown projection | Share of conifers based on crown projection | m^2^ ha^-1^ |  |
| Diameter at breast height (dbh) | Mean diameter at breast height | cm | x |
| Standard deviation of dbh | Standard deviation of diameter at breast height of trees | cm |  |
| Main tree species | Main tree species | deciduous vs. coniferous |  |
| Morisita's index of dispersion | Horizontal dispersion, <1: regular, >1: clumping, 1: random; 20 m × 20 m raster cells |  |  |
| Silvicultural management intensity | Estimated quantitative measure of forest management intensity by Schall and Ammer (2013) |  |  |
| Species diversity | Species diversity based on abundance = inverse Simpson’s index |  |  |
| Stand density | Number of trees | trees ha^-1^ |  |
| Zenner’s Structural Complexity Index | Based on tree height, a proxy for vertical structural complexity |  | x |
| **Geographical variables** |  |  |  |
| Aspect×Slope | Inclination multiplied by 1 for south-, -1 for north-, and 0.5 for east- and west-facing slopes |  | x |
| Exploratory site | Schwäbische Alb: Southwest of Germany, or Hainich-Dün: centre of Germany | Alb and Hainich | x |
| **Microclimatic variables** |  |  |  |
| **Spring (February - May)** |  |  |  |
| Air temperature | Air temperature measured 2 m above ground | °C | x |
| Air temperature (low) | Air temperature measured 10 cm above ground | °C |  |
| Growing days | Days with temperature in the range of 10°C to 30°C | Number of days |  |
| Growth sum | Sum of mean day temperature > 5°C (minus 5) |  |  |
| Warm sum | Sum of mean day temperature with > 10 °C (minus 10) |  |  |
| Soil temperature | Soil temperature at 10 centimeters below surface during winter | °C |  |
| Relative air humidity | Mean relative air humidity measured 2 m above ground during spring | % | x |
| Soil moisture | Soil moisture at 10 centimeters below surface during spring | % |  |
| **Winter (October - January)** |  |  |  |
| Air temperature | Air temperature measured 2 m above ground | °C |  |
| Air temperature (low) | Air temperature measured 10 cm above ground | °C |  |
| Ice days | Days with a temperature maximum < 0°C | Number of days |  |
| Cold days | Days with a temperature minimum < 0°C during winter | Number of days |  |
| Cold sum | The sum of days with a mean day temperature < 0 °C during winter |  |  |
| Cool days | Days with a temperature maximum < 10°C | Number of days |  |
| Soil temperature | Soil temperature at 10 centimeters below surface during winter | °C |  |
| Soil moisture | Soil moisture at 10 centimeters below surface during winter | % |  |
| Relative air humidity | Mean relative air humidity measured 2 m above ground | % |  |
